# Endoplasmic Reticulum Associated Lipolysis Regulates Hepatic Fat Synthesis and Turnover

**DOI:** 10.64898/2026.05.08.723884

**Authors:** Jihong Lian, Russell Watts, Randal Nelson, John P. Kennelly, Aducio Thiesen, Ariel D. Quiroga, Donna Vine, Robin D. Clugston, René L. Jacobs, Richard Lehner

## Abstract

Metabolic Dysfunction-Associated Steatotic Liver Disease (MASLD) is characterized and initiated by the excessive accumulation of triacylglycerols (TG) and cholesteryl esters (CE) in the liver. Hepatic TG and CE synthesis, lipolysis and transport are tightly regulated by nutritional status, and disruption of this homeostasis contributes to MASLD pathogenesis. We have found that an endoplasmic reticulum-localized arylacetamide deacetylase (AADAC) catalyzes hepatic TG/CE turnover, and suppresses SREBP- and LXR-regulated lipogenesis and fatty acid esterification. Consequently, AADAC deficiency in mice leads to increased hepatic lipid synthesis, exacerbated steatosis, and impaired whole-body metabolism during Western-type diet feeding. These findings implicate AADAC as an important regulator of hepatic neutral lipid metabolism, linking endoplasmic reticulum cholesteryl ester hydrolysis as a modulator of lipid synthesis, and suggest its potential role in limiting MASLD pathogenesis under conditions of chronic overnutrition.

## Introduction

Metabolic Dysfunction-Associated Steatotic Liver Disease (MASLD) affects ∼38% of adults worldwide and is a leading cause of chronic liver disease (*1*). MASLD is characterized and initiated by excess storage of triacylglycerol (TG) and cholesteryl esters (CE) in lipid droplets (LD) (*2*). Various sources of fatty acids (FA) contribute to lipid accumulation during the development of MASLD. During fasting, increased FA flux from adipose tissue to the liver augments TG synthesis (*3*). In the postprandial state, substrates for hepatic TG and CE synthesis are provided via uptake of chylomicron remnants and by increased FA and cholesterol syntheses regulated through activation of sterol regulatory element-binding proteins (SREBP) 1c and 2 transcriptional activities, respectively (*4*). SREBP2 activation is tightly regulated by changes in endoplasmic reticulum (ER) cholesterol concentration (*5*). SREBPs thus control the synthesis of lipids required for cellular functions and energy storage, including FA, TG, cholesterol and phospholipids. Metabolic disorders, particularly hyperinsulinemia, lead to elevated hepatic SREBP1c activation, making *de novo* lipid synthesis a significant contributor to the development of MASLD (*6*).

Conversely, hepatic TG and CE mobilization and clearance depend on lipolysis as a key step. Lipases cleave ester bonds in TG and CE, releasing FA, glycerol and cholesterol for various downstream metabolic processes. The main lipases catalyzing these processes in the liver include adipose triglyceride lipase (ATGL) (*7*), hydrolyzing TG stored in cytosolic LDs, and lysosomal acid lipase (LAL), hydrolyzing TG and CE present in endocytosed chylomicron remnants and in LDs via lipophagy (*8*). During fasting, FA released by ATGL and lipophagy are delivered to mitochondria for oxidation and ketone body production (*7, 8*). Provision of substrates for very low-density lipoprotein (VLDL) assembly in the ER also requires lipolysis of preformed TG stores (*9*) and this process is catalyzed by the ER-localized carboxylesterase Ces1d in mice (*10*) and its ortholog CES1 in humans (*11*). Hepatic neutral-pH CE hydrolase(s) mobilizing CE from LD remain poorly defined. Hormone-sensitive lipase (HSL) is a known CE hydrolase that also hydrolyzes diacylglycerol, TG, and other esters (*12*). However, HSL is weakly expressed in mouse hepatocytes compared to immune cells in the liver (*13*) and is absent from human hepatocytes (*14*), making its role in hepatocyte lipid turnover unclear and unlikely to be major. Given the importance of neutral lipid mobilization and the pathological consequence of its dysregulation, characterization of hepatic lipolysis under different nutritional states is essential.

Arylacetamide deacetylase (AADAC) was initially isolated from liver intracellular membrane preparations and characterized as a drug-detoxifying enzyme *in vitro* (*15*). AADAC is an ER membrane glycoprotein with the active site oriented towards the ER lumen (*16*). Interestingly, AADAC was identified as a LD-associated protein in mouse liver (*17*). SAY1, the yeast ortholog of AADAC, was also present in the yeast LD proteome These findings suggested that AADAC could be localized at ER-LD contact sites The catalytic site of AADAC shares homology with HSL including the conserved oxyanion forming residues (H_110_GGG), the serine lipase/hydrolase active site motif (GXS_188_XG), and lipase/esterase catalytic triad residues (S_188_D_342_H_372_) (*20*). Unlike HSL, AADAC is predominantly expressed in hepatocytes in both mouse (*13*) and human (*14*) liver. We and others have demonstrated that AADAC exhibits lipase activities towards acylglycerols and steryl esters (*16, 21*). In line with these studies, overexpression of AADAC in hepatoma cells (*16*) or vascular smooth muscle cells (*22*) decreased cellular lipid storage.

Based on the intracellular localization and lipolytic activity of AADAC, we hypothesized that AADAC activity may contribute to neutral lipid turnover in hepatocytes and influence *de novo* lipid synthesis by modulating SREBP activation. Mutations in the human *AADAC* gene that may affect AADAC abundance and/or activity have been reported (*23, 24*) but whether these mutations are associated with altered lipid metabolism is unknown. Knockdown of endogenous *AADAC* in a human liver cell line Huh7.5 decreased lipolysis of stored TG (*25*). Furthermore, AADAC activity was found to be decreased in the liver of obese individuals (*26*). *AADAC* expression is downregulated in metabolic dysfunction-associated steatohepatitis (MASH) (*27*). To address the role of AADAC in lipid metabolism *in vivo* we generated global AADAC knockout (KO) mice. Our data show that AADAC deficiency blunts mobilization of both TG and CE during the transition from fasting to refeeding in the liver and increases *de novo* lipid synthesis through upregulation of SREBP and LXR pathways, leading to exacerbated MASLD pathology.

## Results

### Tissue distribution and expression pattern of AADAC protein

To interrogate the physiological function of AADAC in whole body metabolism, we generated AADAC KO mice (fig. S1A). AADAC deficiency decreased total esterase activity in the mouse liver homogenate and microsomal fraction (fig. S1B).

AADAC protein is most abundant in liver, with much lower expression in the small intestine and kidney (fig. S1C), and was not detected in other tissues suggesting either absence or extremely low abundance.

The expression of AADAC protein and mRNA were decreased by 37% and 26%, respectively, in the livers of leptin-deficient, obese, insulin resistant *ob/ob* mice (fig. S1D). HSL hydrolyzes retinyl ester (RE) to retinol (*28*). Given the active site homology between AADAC and HSL, we examined whether AADAC deficiency influences hepatic RE metabolism. Liver RE and retinol levels were comparable between genotypes (fig. S1E), suggesting that AADAC does not play an important role in hepatic RE hydrolysis *in vivo*.

### AADAC deficiency leads to liver lipid accumulation during short-term Western-type diet feeding

To assess whether AADAC deficiency affects lipid homeostasis during nutritional challenge, we fed both WT and AADAC KO male and female mice a high-fat, high-cholesterol, high-sucrose Western-type diet (WTD) for one week. Liver lipid profiles were measured in fasted, and 2-hour and 6-hour refed states. Male WT mice exhibited significant hepatic TG and CE turnover during the transition from fasted to refed state, whereas this turnover was blunted in the livers of AADAC KO mice, leading to increased postprandial hepatic TG and CE concentrations (Fig. 1A). FC levels were not altered by feeding conditions, but KO mice showed a slight increase in FC concentration in the 2-hour refed state compared with WT mice (Fig. 1A). These data suggested that AADAC is responsible for liver TG and CE turnover particularly during the prandial state.

**Fig. 1.**
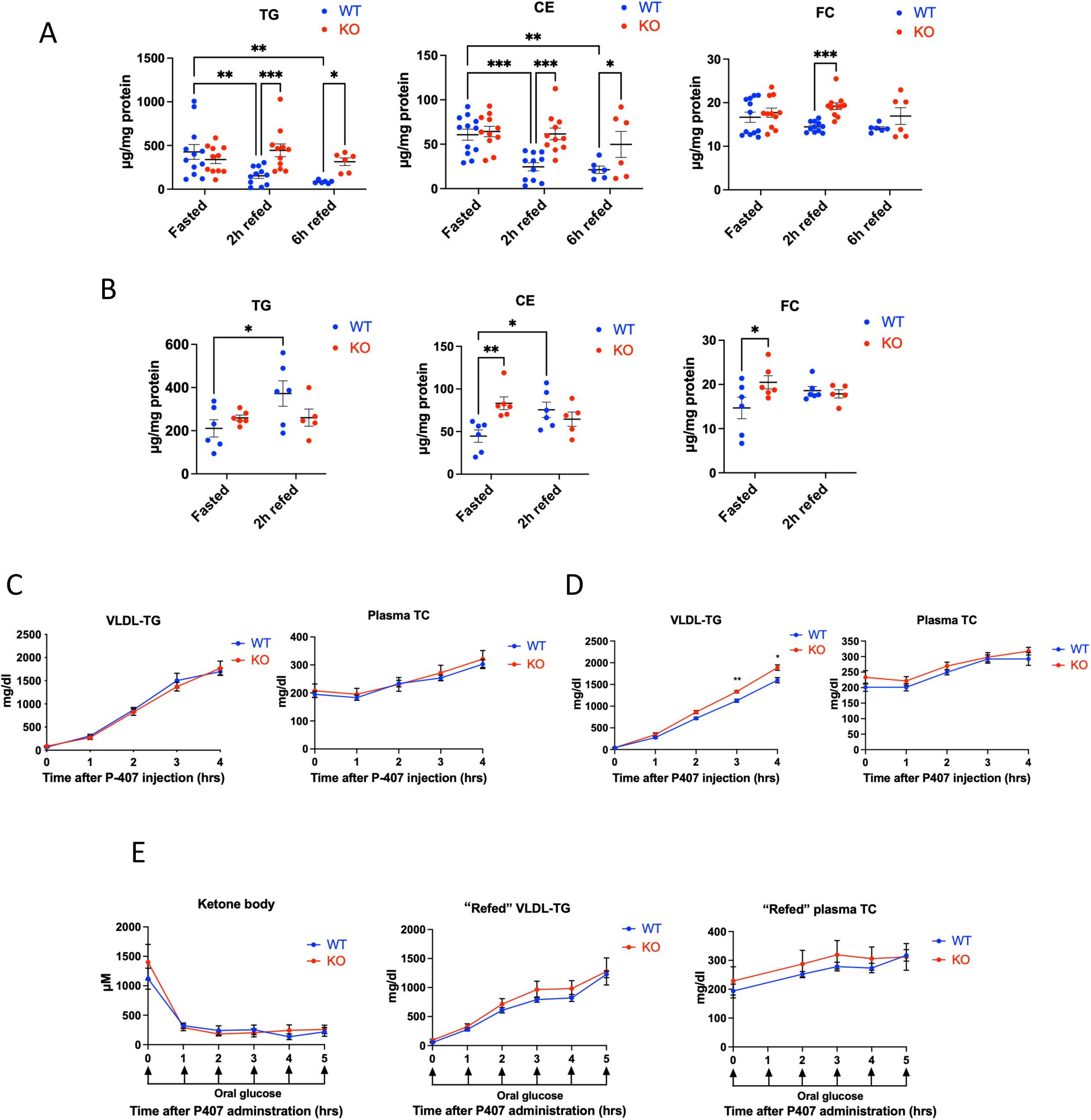
AADAC deficiency leads to hepatic neutral lipid accumulation in mice fed Western-type diet (WTD) for one week. **(A)** Liver triacylglycerol (TG), cholesteryl ester (CE), and free cholesterol (FC) in fasted and refed male WT and AADAC KO mice fed WTD for one week. **(B)** Liver TG, CE, and FC in fasted and refed female WT and AADAC KO mice fed WTD for one week. **(C)** VLDL-TG and total cholesterol (TC) secretion in fasted male WT and AADAC KO mice fed WTD for one week, n=6. **(D)** VLDL-TG and TC secretion in fasted female WT and AADAC KO mice fed WTD for one week, n=6. **(E)** Ketone body concentrations, and VLDL-TG and TC secretion in male WT and AADAC KO mice fed WTD for one week were assessed under a condition to mimic fed state by giving the mice oral glucose every hour following overnight fasting and P-407 administration, n=5. Data are expressed as mean ± SEM. *P<0.05, **P<0.01, ***P<0.001. Statistical significance was determined by two-way ANOVA followed by Bonferroni post hoc tests.

Unlike male WT mice, female WT mice exhibited lower CE and TG levels in the fasted state than the refed state (Fig. 1B), potentially due to sex differences in systemic FA partitioning (*29*). However, female AADAC KO mice still had increased hepatic CE levels during fasting compared to WT mice, and the differences in neutral lipid levels between fasting and refeeding were absent in the female KO livers (Fig. 1B).

### The accumulation of hepatic lipids in WTD fed AADAC deficient mice is not due to attenuated VLDL secretion

ER-associated lipolysis plays a crucial role in providing lipid substrates for VLDL assembly (*9*). To investigate whether the accumulation of liver lipids in the WTD fed AADAC KO mice results from decreased VLDL secretion, we performed *in vivo* VLDL secretion tests in both fasted and refed conditions. After one week of WTD feeding, male AADAC KO mice showed no difference in fasted VLDL-TG secretion and plasma TC concentration (Fig. 1C). Female AADAC KO mice exhibited increased VLDL-TG secretion but did not show a significant difference in plasma TC concentration (Fig. 1D).

To address the VLDL secretion rate during refeeding in male WT and AADAC KO mice, we performed the VLDL secretion assay under a condition mimicking the fed state by giving mice oral glucose every hour following overnight fasting—this approach avoids the promotion of bulk lipid secretion from the intestine. Ketone body concentration was dramatically supressed after oral glucose administration, indicating the feeding response was stimulated in these mice (Fig. 1E). VLDL-TG secretion and plasma TC were not different between WT and KO animals in the fed state (Fig. 1E). These results suggest that the difference observed in liver lipids is not due to decreased lipid secretion via VLDL from the liver.

### AADAC deficiency is not associated with marked changes in intestinal lipid metabolism

Given that the intestine is the second major site of AADAC expression, we examined the role of AADAC in intestinal lipid metabolism. No significant changes in TG or TC in either male or female KO mice compared to WT mice were observed in jejunal mucosa collected after 2-hour refeeding following one week of WTD (fig. S2A), consistent with histological examination showing no detectable differences in intestinal lipid accumulation (fig. S2B).

Microarray analysis of intestinal mucosa from male mice fed one week of WTD identified 42 and 65 differentially expressed genes (DEGs) in the fasted and 2-hour refed states, respectively, in KO relative to WT mice (fig. S2C, and table S1, A and B), suggesting that loss of AADAC does not induce major transcriptional remodeling in intestinal mucosa. Notably, none of these DEGs were canonical regulators of intestinal lipid metabolism. Gene Ontology (GO) biological process and KEGG pathway analyses revealed only a limited number of significantly enriched terms, none of which were related to lipid metabolic pathways (table S1, C to G).

Plasma TG and TC levels after 2-hour refeeding were comparable between male WT and KO mice (fig. S2D), indicating that postprandial blood lipid levels, which are largely derived from intestinal chylomicron secretion, were not influenced by AADAC deficiency, and suggesting that the increased hepatic lipid levels observed in refed KO mice are unlikely to result from altered chylomicron remnant delivery to the liver.

### AADAC deficiency accelerates MASLD progression and modestly impairs whole-body metabolism after long-term WTD feeding

The WTD-fed mouse is a dietary model that closely recapitulates human MASLD both metabolically and histologically (*30*). Both male WT and AADAC KO mice developed severe steatosis after 10-week WTD feeding, as determined by pathohistological scoring (Fig. 2A) and lipid profiles (Fig. 2B). Although liver TG concentration in male AADAC KO mice was not significantly different from WT mice, AADAC KO mice exhibited significantly increased liver CE (Fig. 2B). Female WT mice developed milder steatosis than male WT mice (Fig. 2C), and female AADAC KO mice showed greater pathohistological scoring of MASLD (Fig.2C), and increased liver TG and CE concentrations (Fig. 2D) compared to WT mice.

**Fig. 2.**
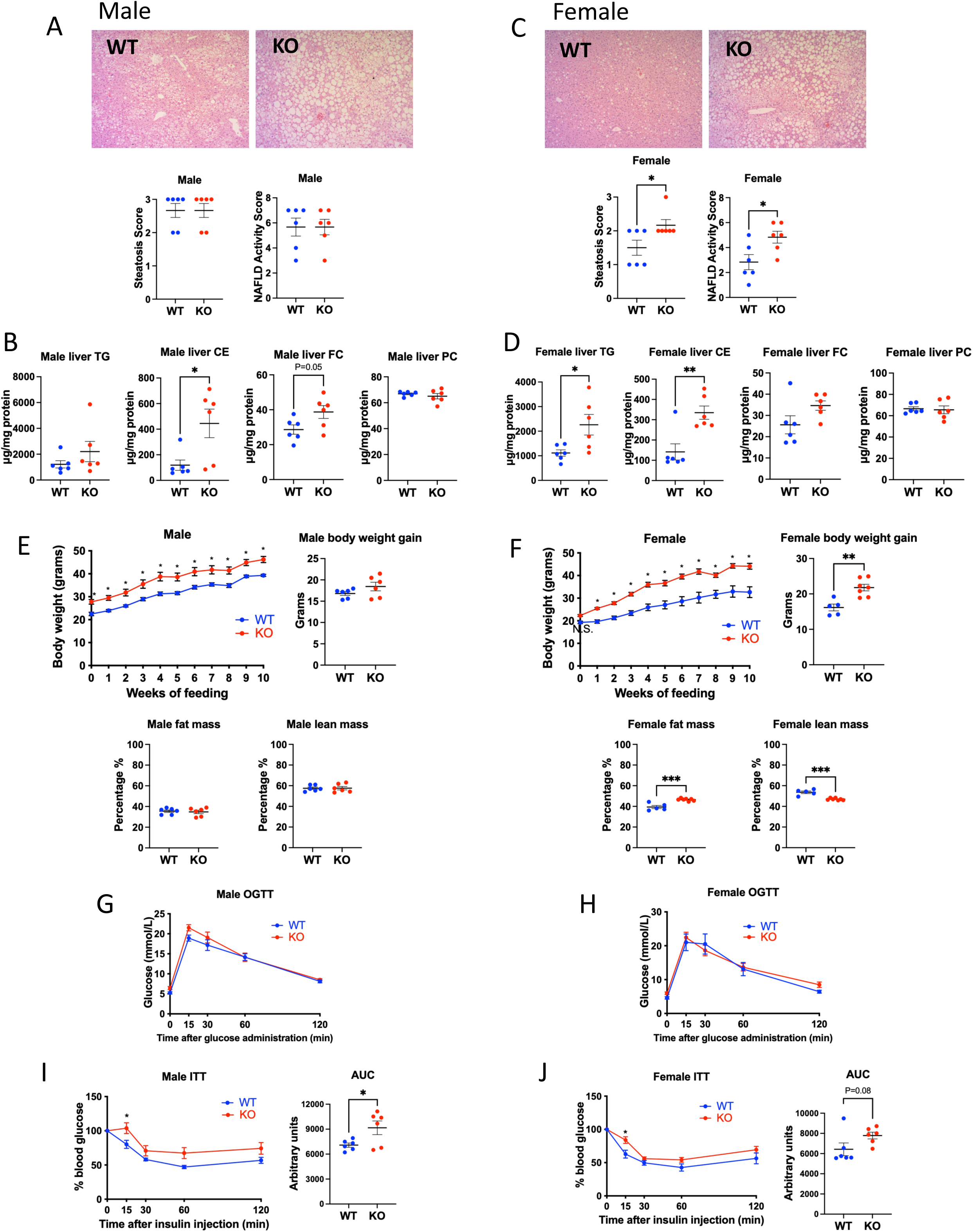
AADAC deficiency exacerbates WTD-induced MASLD development and affects whole body metabolism. **(A)** Histopathological analysis of MASLD in male WT and AADAC KO mice fed WTD for 10 weeks. Steatosis score (scale: 1-3) and NAFLD (MASLD) activity score were quantified. **(B)** Liver lipid profile of male WT and AADAC KO mice fed WTD for 10 weeks. PC, phosphatidylcholine. **(C)** Histopathological analysis of MASLD in female WT and AADAC KO mice fed WTD for 10 weeks. Steatosis score (scale: 1-3) and NAFLD (MASLD) activity score were quantified. **(D)** Liver lipid profile of female WT and AADAC KO mice fed WTD for 10 weeks. Body weight, body weight gain, and body composition of **(E)** male and **(F)** female WT and AADAC KO mice fed WTD for 10 weeks, n=6. Oral glucose tolerance test (OGTT) in **(G)** male and **(H)** female WT and AADAC KO mice fed WTD for 10 weeks, n=6. Insulin tolerance test (ITT) in **(I)** male and **(J)** female WT and AADAC KO mice fed WTD for 10 weeks, n=6. Data are expressed as mean ± SEM. *P<0.05, **P<0.01, ***P<0.001. Statistical significance was determined by two-way ANOVA followed by Bonferroni post hoc tests, or by unpaired two-tailed t-tests for two-group comparisons.

MASH is distinguished from simple steatosis by inflammation, hepatocyte injury and fibrosis (*31*). In male AADAC KO mice, 10-week WTD increased hepatic expression of the macrophage marker *Cd68* and the proinflammatory cytokine *Ccl2* compared to WT mice on the same diet or chow fed controls (fig. S3A). In addition, mRNA abundance of genes encoding the macrophage marker F4/80 and the proinflammatory cytokine Tnfα were increased by WTD only in KO livers. Female KO mice showed elevated *Ccl2* mRNA abundance after 10-week WTD feeding (fig. S3B). The mRNA abundance of *Col1a1* encoding fibrosis marker collagen type 1 increased in both sexes of AADAC KO mice after 10-week WTD feeding (fig. S3, A and B). These data indicate exacerbated diet-induced MASH features in AADAC KO mice of both sexes.

At baseline, male AADAC KO mice were heavier than WT mice while no difference was observed in female mice (Fig. 2, E and F). During the 10-week WTD feeding, male KO mice remained heavier than WT mice without differences in weight gain (Fig. 2E), while female KO mice gained more weight than WT mice (Fig. 2F). Consistently, fat and lean mass percentages were comparable between male WT and KO mice (Fig. 2E), whereas female KO mice exhibited increased fat mass and decreased lean mass percentage (Fig. 2F). The decreased lean mass percentage in female KO mice was not due to absolute lean mass loss (fig. S3C).

While both male and female AADAC KO mice did not show differences in glucose tolerance after long-term WTD feeding (Fig. 2, G and H), the AADAC KO mice exhibited more pronounced insulin resistance (Fig. 2, I and J).

### Impaired lipolysis in AADAC deficient hepatocytes

We interrogated the mechanisms by which AADAC affects lipid metabolism in hepatocytes isolated from WT and KO mice after 12-hour fasting, 2-hour refeeding, and 6-hour refeeding. AADAC deficient hepatocytes exhibited decreased LD number (Fig. 3, A and B) and increased LD size (Fig. 3, A and C) under fasting and 2-hour refeeding conditions when compared with WT hepatocytes. In hepatocytes isolated after 6-hour refeeding, large LDs were nearly depleted in WT but retained in KO hepatocytes (Fig. 3, A and C), suggesting impaired neutral lipid turnover.

**Fig. 3.**
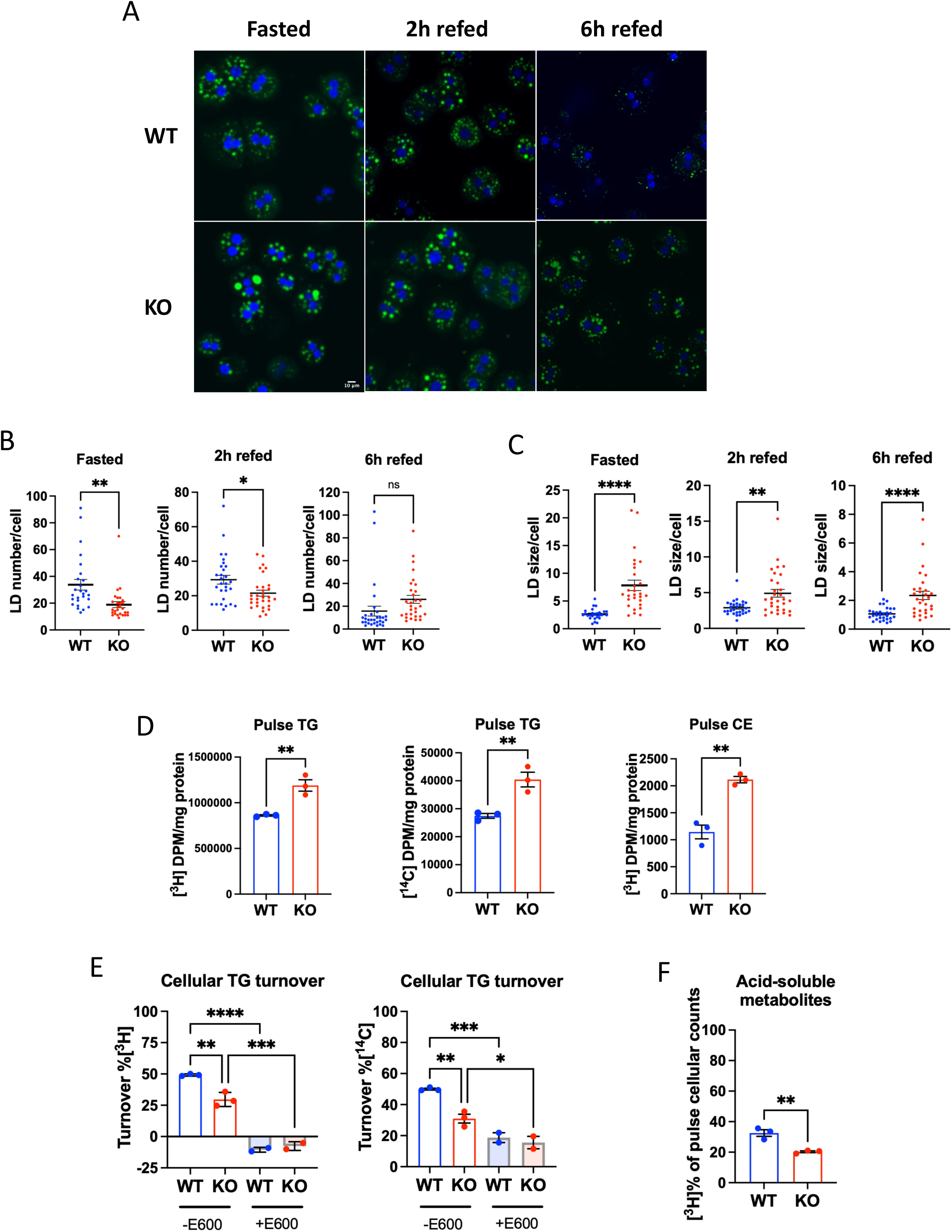
Impaired lipolysis in AADAC deficient primary hepatocytes. **(A)** LDs were visualized by BODIPY 493/503 staining, and **(B)** LD numbers and **(C)** sizes were analyzed in hepatocytes isolated from fasted and refed male WT and AADAC KO mice. **(D)** Pulse-chase experiment was performed to assess turnover of TG in WT and AADAC KO hepatocytes. TG was labeled by [^3^H]oleic acid (OA) and [^14^C]glycerol in the pulse phase, incorporation of FA into CE in the pulse phase was also analyzed. **(E)** Turnover of preformed cellular TG was assessed in the chase phase in the presence of DGAT inhibitors and in the presence or absence of general lipase inhibitor E600. **(F)** Release of lipolytic metabolites (acid-soluble metabolites) in the chase phase was measured in the culture medium. Data are expressed as mean ± SEM. Ns, not significant, *P<0.05, **P<0.01, ***P<0.001, ****P<0.0001. Statistical significance was determined by one-way ANOVA followed by Bonferroni post hoc tests, or by unpaired two-tailed *t*-tests for two-group comparisons.

To test our hypothesis that the sustained large LDs in AADAC KO hepatocytes resulted from impaired lipolysis we performed a radioactive double-label pulse-chase experiment in primary hepatocytes. To mimic neutral lipid synthesis from exogenous FAs during fasting, hepatocytes isolated from fasted WT and KO mice were incubated in medium containing fasted human serum, [^3^H]oleic acid (OA) and [^14^C]glycerol used to label FAs and the glycerol backbone of TG in newly synthesized lipids, respectively. After 4 hours of labeling (pulse period), increased incorporation of [³H]OA and [¹⁴C]glycerol into TG, as well as increased [³H]OA incorporation into CE, were observed in KO hepatocytes (Fig. 3D). To investigate the role of AADAC in the hydrolysis of stored lipids, we analyzed turnover of preformed labeled lipids in WT and KO hepatocytes in the presence of insulin to mimic the refeeding response. DGAT1/2 inhibitors were included to prevent re-esterification of released FA back to TG. About 50% of pulse-labeled [^3^H]- and [^14^C]-TG was turned over in WT hepatocytes compared to 20% in KO (Fig. 3E). Low turnover was observed in cells treated with the pan lipase inhibitor E600, indicating that some of the decreased lipid turnover in KO hepatocytes was due to reduced lipase activity (Fig. 3E). In agreement with the decreased cellular TG turnover in KO hepatocytes, we also observed decreased production of acid-soluble metabolites in the medium, suggesting diminished FA release and availability for oxidation (Fig. 3F).

### Accumulation of lipids in the liver of AADAC deficient mice is independent of PLIN5-ATGL regulated lipolysis

We investigated the impact of other major regulators of hepatic lipid storage and lipolysis in WTD fed AADAC KO mice. Perilipins (PLINs) are well-characterized LD coat proteins. PLIN2 is a constitutive LD coat protein, and its abundance correlates with lipid storage in the liver (*32*). PLIN2 abundance was significantly increased in male AADAC KO livers during fed state compared to WT livers (Fig. 4A). PLIN5, which interacts with ATGL and ATGL activator ABHD5/CGI-58 and thereby regulates TG lipolysis in LDs (*33*), was also increased in the liver of AADAC KO mice (Fig. 4A). Interestingly, increased TG storage in AADAC KO livers during feeding was observed despite augmented ATGL abundance (Fig. 4A). To investigate whether increased PLIN5 blocked ATGL-mediated TG hydrolysis in the liver of AADAC KO mice, thereby contributing to blunted lipid turnover and increased TG accumulation in the fed states, we attenuated PLIN5 expression in AADAC KO mice using a liver-targeted PLIN5 antisense oligonucleotide (ASO) (Fig. 4B). Hepatic lipid concentrations remained unchanged between PLIN5 ASO- and control ASO-treated KO mice (Fig. 4C), indicating lipid accumulation, especially TG accumulation in the AADAC KO livers was not due to blockage of lipolysis by increased PLIN5 abundance. The increased hepatic TG in PLIN5 ASO-treated WT mice aligns with findings from liver-specific PLIN5 KO mice, which developed exacerbated steatosis (*34*). The elevated PLIN5 levels in KO livers may be proportional to the increased LD content. Consistent with this hypothesis, there was no difference in the abundance of PLIN5 and ATGL in the livers of female KO mice in the fed state after one week of WTD feeding (fig. S4), which also exhibited no differences in lipid levels under this condition.

**Fig. 4.**
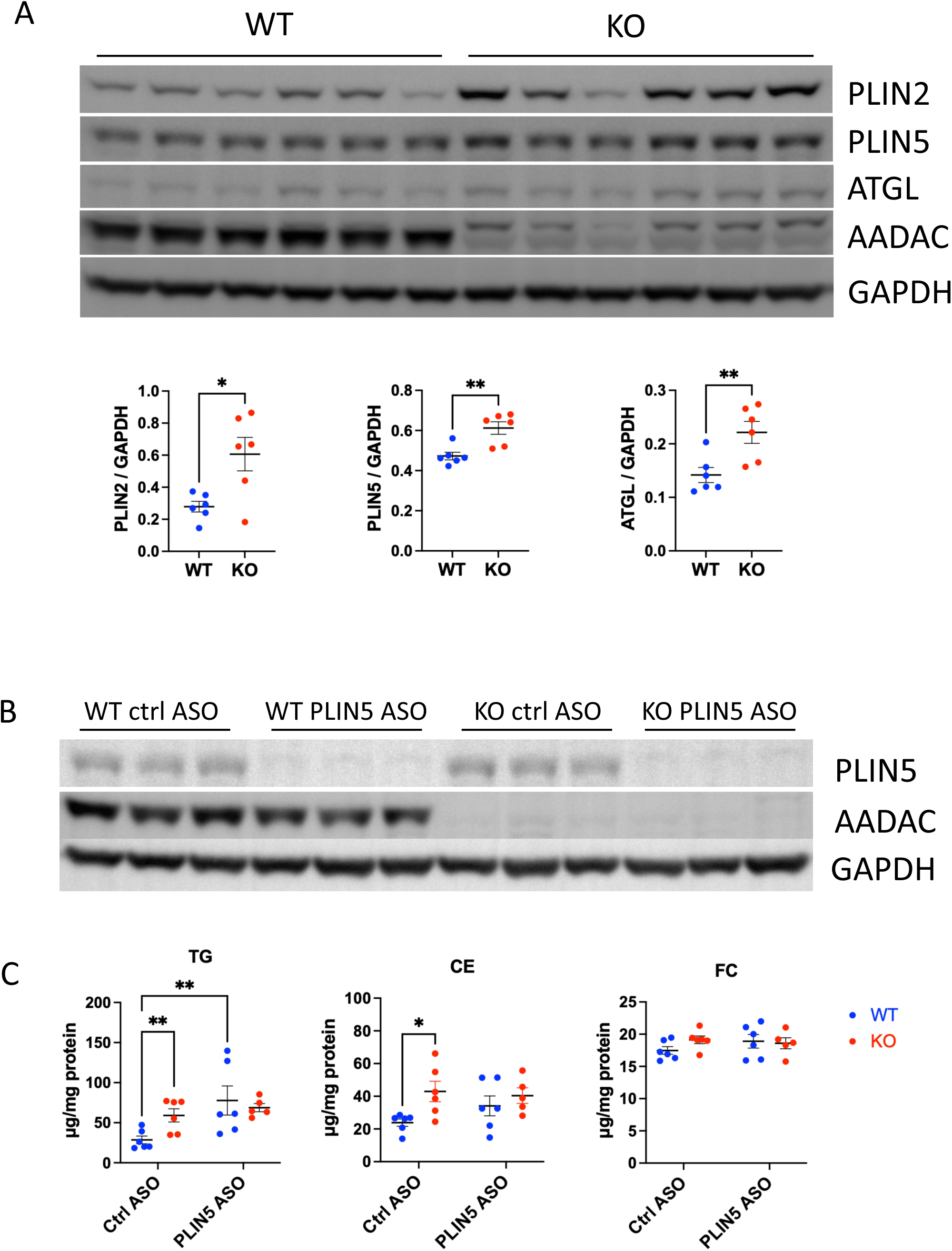
Lipid accumulation in the livers of AADAC deficient mice is independent of PLIN5-ATGL regulated lipolysis. **(A)** Liver protein abundance of PLIN2, PLIN5, and ATGL in male WT and AADAC KO mice in the fed state after one week of WTD. **(B)** Hepatic PLIN5 ablation by antisense oligonucleotide (ASO). **(C)** Liver lipid profile in the fed state in ASO-treated mice after one week of WTD feeding. Data are expressed as mean ± SEM. *P<0.05, **P<0.01. Statistical significance was determined by two-way ANOVA followed by Bonferroni post hoc tests, or by unpaired two-tailed *t*-tests for two-group comparisons.

### AADAC deficiency enhances lipid metabolic pathways and leads to increased hepatic *de novo* lipogenesis

To explore the global impact of AADAC deficiency on hepatic lipid and energy metabolism, we performed RNA sequencing (RNAseq) to generate a comprehensive transcriptomic profile in the liver of WT and KO mice collected in the fed state after one week of WTD feeding. Differential expression analysis identified 1,051 DEGs in male AADAC KO mice compared to WT controls (False Discovery Rate [FDR] < 0.05), comprising 592 upregulated and 459 downregulated genes (table S2, A and B), as visualized in the volcano plot (fig. S5, A and B). Numerous genes involved in hepatic lipid metabolism displayed a differential expression pattern between male KO and WT mice (fig. S5C). Fewer DEGs were identified in the livers of female mice (26 upregulated and 20 downregulated genes) (table S2, C and D).

GO analysis, particularly the biological process category (table S2G), and the KEGG pathway analysis (table S2I) identified multiple lipid metabolism pathways that were significantly upregulated in the liver of male AADAC KO mice (Fig. 5A). Notably, biological processes including cholesterol biosynthetic process (GO:0006695), fatty acid biosynthetic process (GO:0006633), triglyceride biosynthetic process (GO:0019432), and lipid transport (GO:0006869), were overrepresented among upregulated DEGs in KO mice (Fig. 5, A and B). The upregulated DEGs within these enriched pathways include the SREBP2 target genes *Insig1*, *Dhcr24*, and *Ldlr* (Fig. 5B), suggesting increased SREBP2 activation. Upregulation of the *Srebf1* gene was also observed in the KO liver. Multiple SREBP1c target genes, including those encoding key lipogenic enzymes, *Fasn*, *Scd1*, and *Acaca* (encoding ACC1), as well as other lipid synthesis-related genes (*Gpam* encoding GPAT1, *Acsl5, Elovl5, and Elovl6*), were upregulated in the KO livers (Fig. 5B). Within the biological process of “lipid transport”, multiple canonical liver X receptor (LXR) target genes, including *Abcg5, Abcg8, Abca1*, and *Cd36*, were upregulated in the KO livers (Fig. 5B), suggesting increased LXR activation. LXR is known to transcriptionally upregulate the *Srebf1* expression (*35*). Additionally, LXR binds to an LXR response element within the promoter regions of key lipogenic genes such as *Fasn* and *Acaca* and enhances their transcription (*36, 37*). Because SREBP2 has been demonstrated to increase *Srebf1* gene transcription through activating LXR (*38*), the overall upregulation of these lipid metabolism-related genes in AADAC KO livers suggests a mechanistic pathway driving increased lipid synthesis. In this pathway, activated SREBP2 upregulates *Srebf1* expression and its target genes through LXR activation, while LXR can also directly enhance lipogenic gene expression.

**Fig. 5.**
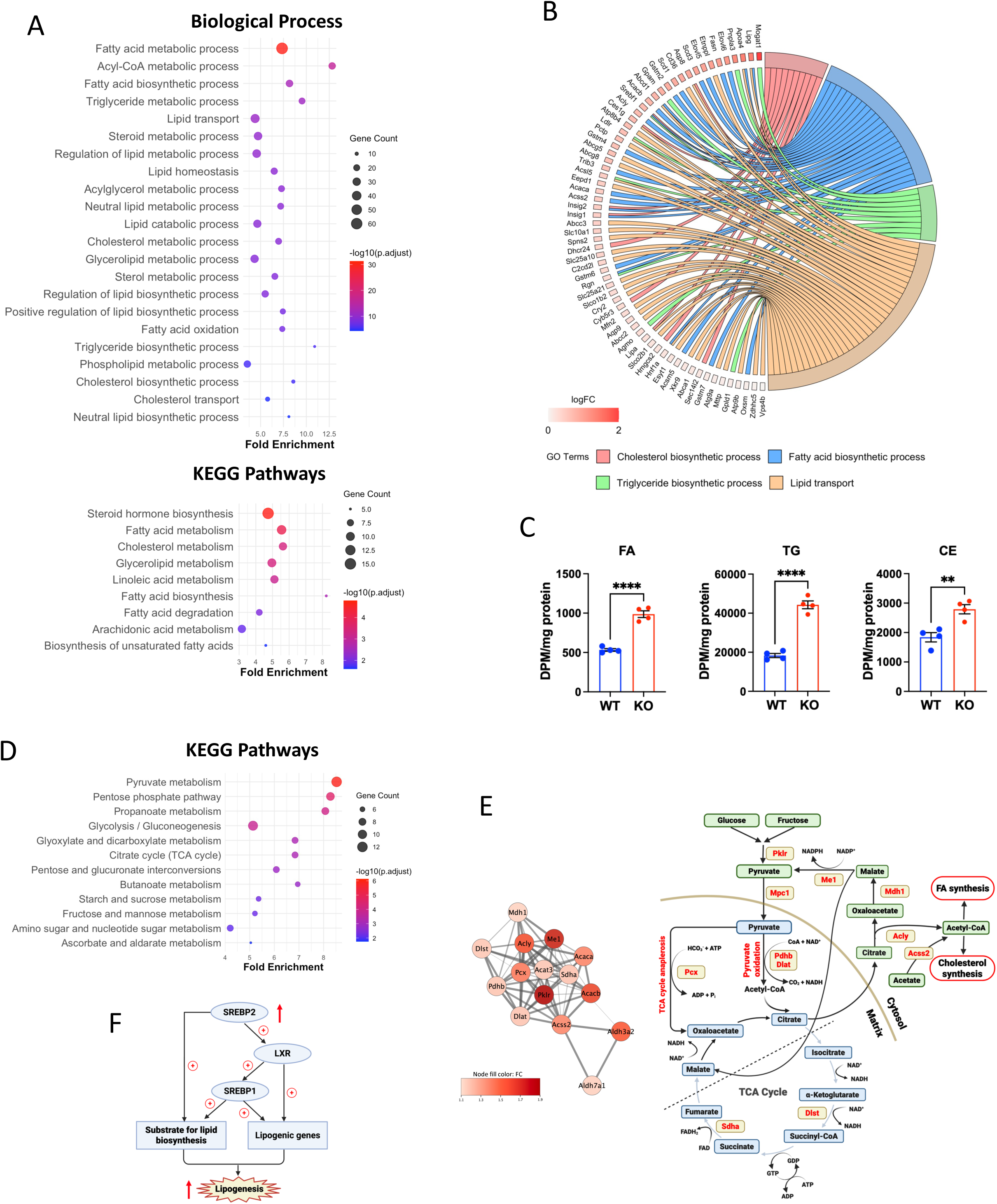
Upregulation of lipid biosynthesis pathways and increased *de novo* lipogenesis in the livers of AADAC deficient mice. **(A)** Upregulated biological processes and KEGG pathways in lipid metabolism in the livers of male AADAC KO mice after one week of WTD feeding. The top enriched biological processes and pathways ranked by adjusted p-value are shown. **(B)** Upregulated differentially expressed genes (DEGs) in the biological processes of cholesterol biosynthetic process, fatty acid biosynthetic process, triglyceride biosynthetic process, and lipid transport in the livers of male AADAC KO mice after one week of WTD feeding. (C) *De novo* lipogenesis was assessed by measuring incorporation of [^3^H]acetic acid into FA, TG, and CE in primary hepatocytes isolated from WT and AADAC KO mice. Data are expressed as mean ± SEM. **P<0.01, ****P<0.0001. Statistical significance was determined by unpaired two-tailed *t*-tests. **(D)** Upregulated KEGG pathways related to carbohydrate metabolism in the livers of male AADAC KO mice after one week of WTD feeding. The top enriched pathways ranked by adjusted p-value are shown. **(E)** Remodeling of TCA cycle and related pathways in the livers of male AADAC KO mice. Left panel: STRING network of DEGs in enriched pyruvate metabolism and citrate cycle (TCA cycle) pathways. Edge thickness corresponds to interaction confidence scores from STRING; node color intensity reflects the magnitude of fold change. Right panel: mapping of upregulated DEGs (KO vs WT) in the TCA cycle and related pathways. *Acly*, ATP citrate lyase; *Acss2*, acetyl-CoA synthetase 2; *Dlat*, dihydrolipoamide S-acetyltransferase; *Dlst*, dihydrolipoamide S-succinyltransferase; *Mdh1*, malate dehydrogenase 1; *Me1*, malic enzyme 1; *Mpc1*, mitochondrial pyruvate carrier 1; *Pcx*, pyruvate carboxylase; *Pdhb*, pyruvate dehydrogenase beta; *Pklr*, pyruvate kinase L/R; *Sdha*, succinate dehydrogenase complex subunit A. **(F)** Cascade leading to increased *de novo* lipogenesis in the livers of AADAC KO mice after one week of WTD feeding.

While female KO and WT livers showed limited gene expression differences, female WT livers displayed dramatic upregulation of lipid biosynthetic processes compared to male WT livers (table S2K). The same four typical biological processes upregulated in male KO livers (Fig. 5B) were all enriched among upregulated DEGs in female WT versus male WT comparisons (fig. S5D, and table S2E). This constitutively elevated expression of lipid biosynthetic genes in female livers may explain their reduced sensitivity to the KO condition.

We investigated if *de novo* lipogenesis is functionally affected by AADAC deficiency. Hepatocytes were isolated from WT and AADAC KO mice during the fed state when *de novo* lipogenesis is elevated, and acetate labeling was performed in the presence of human refed serum. We observed increased incorporation of the labelled precursor into FA and neutral lipids in AADAC KO hepatocytes (Fig. 5C), indicating increased lipogenesis.

### AADAC deficiency enhances pathways providing substrates for *de novo* lipid synthesis

KEGG pathway analysis revealed multiple carbohydrate metabolism-related pathways among upregulated DEGs in the KO livers, including pyruvate metabolism (mmu00620) and citrate cycle (TCA cycle; mmu00020) (Fig. 5D). STRING network analysis followed by MCL clustering of the 15 DEGs from these two pathways identified an interconnected functional module (Fig. 5E, left panel). *Acly* (encoding ATP citrate lyase) and *Acss2* (encoding acetyl-CoA synthetase 2) emerged as the most highly connected nodes in the network (degree of 12 out of 15 genes), suggesting that they serve as central hubs coordinating metabolic responses to AADAC deficiency in this module, with both enzymes producing cytosolic acetyl-CoA, the key precursor for *de novo* lipid synthesis.

Mapping DEGs in this module onto TCA cycle and related metabolic pathways revealed coordinated upregulation across multiple interconnected routes, supporting a model in which TCA cycle-derived metabolites are redirected toward *de novo* lipid synthesis (Fig. 5E, right panel). Upregulated genes included those involved in pyruvate formation from glycolysis/fructolysis (*Pklr*), pyruvate mitochondrial import (*Mpc1*), acetyl-CoA formation and citrate-malate shuttle promoting transfer of acetyl-CoA equivalents from mitochondria to cytosol (*Pdhb, Dlat*, *Acly, Mdh1, Acss2*), NADPH production (*Me1*), and TCA cycle maintenance (*Pcx, Dlst, Sdha*). The pentose phosphate pathway (mmu00030), another major NADPH source, was also upregulated in AADAC KO liver (Fig. 5D). Notably, *Acly, Mdh1, Me1* and *Acss2* are all established SREBP targets (*39–42*). These findings suggest that AADAC deficiency promotes metabolic reprogramming to enhance substrate supply for hepatic lipid synthesis through elevated SREBP activity (Fig. 5F).

Concurrently, enrichment of the fructose and mannose metabolism pathway (mmu00051) (Fig. 5D, and table S2I) revealed upregulation of fructolytic genes (*Khk*, *Aldob*, *Tkfc)* in AADAC KO livers, which constitute a complete fructolysis pathway that bypasses key regulatory steps of glycolysis, enabling rapid conversion of fructose derived from sucrose in the WTD into lipogenic precursors.

### Mitochondrial functional remodeling in the liver of AADAC KO mice

Morphological analysis showed more mitochondria surrounding LDs in liver tissue (Fig. 6A) and primary hepatocytes (Fig. 6B) of AADAC KO mice compared to WT controls.

**Fig. 6.**
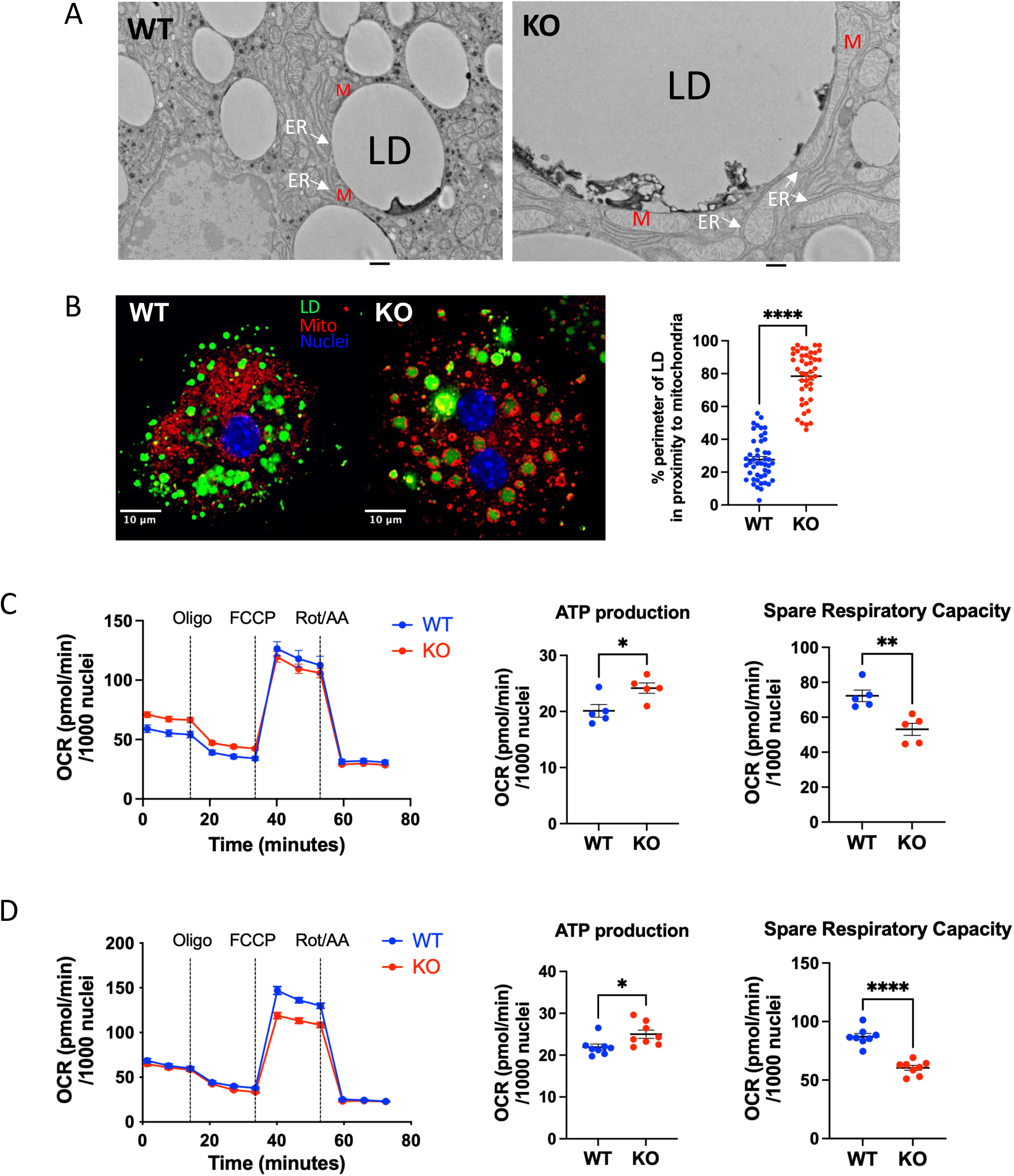
Mitochondrial functional remodeling in the livers of AADAC deficient mice. **(A)** Increased peridroplet mitochondria in the livers of AADAC KO mice, M, mitochondria; LD, lipid droplets; ER, endoplasmic reticulum; scale bar = 500 nm; and **(B)** in primary hepatocytes of AADAC KO mice, green, lipid droplets; red, mitochondria; blue, nucleus; scale bar = 10 μm. **(C)** Seahorse XF mitochondrial stress analysis performed on primary hepatocytes isolated from WT and KO mice fed with chow diet, or **(D)** one week of WTD. Data show mitochondrial stress test kinetics with sequential addition of oligomycin, FCCP, and rotenone/antimycin A (left panels), along with quantified ATP-linked respiration (middle panels), and spare respiratory capacity (SRC, right panels). Data are expressed as mean ± SEM. *P<0.05, **P<0.01, ****P<0.0001. Statistical significance was determined by unpaired two-tailed *t*-tests.

LD-mitochondria interactions have been reported across various cell types (*43–45*), with studies demonstrating that peridroplet mitochondria support lipid synthesis and LD expansion by providing ATP (*43*), and by promoting lipogenesis from non-lipid precursors (*44, 45*). The spatial proximity between these organelles may play a crucial role (*45*).

To investigate the hepatic mitochondrial bioenergetics in AADAC KO mice, Seahorse XF mitochondrial stress analysis was performed on primary hepatocytes isolated from WT and KO mice fed either chow diet or WTD for one week. AADAC deficient hepatocytes exhibited increased ATP-linked respiration (Fig. 6, C and D), suggesting enhanced basal ATP production to meet higher metabolic demands such as lipid biosynthesis. This finding is consistent with increased TCA cycle enzyme expression in AADAC KO livers observed by RNA-seq analysis (Fig. 5E).

Spare respiratory capacity (SRC) represents the reserve capacity of mitochondria to increase oxygen consumption and ATP production above basal levels in response to bioenergetic stress. Notably, SRC is decreased in AADAC KO hepatocytes under both chow diet and WTD fed conditions (Fig. 6, C and D). The mitochondrial marker VDAC1, as well as oxidative phosphorylation (OXPHOS) complexes, showed comparable abundance between WT and KO livers (fig. S6), suggesting the reduced SRC was not due to decreased mitochondrial mass or respiratory chain content. Another known factor influencing SRC is the total availability of mitochondrial substrate that can be oxidized by the TCA cycle (*46*). The reduced SRC in AADAC KO hepatocytes may reflect substrate limitation caused by the reprogramming of the TCA cycle and related metabolic pathways, which diverts carbon flux towards cytosolic lipid anabolism rather than mitochondrial oxidative metabolism (Fig. 5E).

### Structural insights and regulatory mechanisms of AADAC activity

AADAC is an ER-localized type II membrane protein (*16*) with a single transmembrane domain (TMD). The protein structure of mouse AADAC was predicted by AlphaFold 3 (Fig. 7A). The catalytic site is buried in the interior of AADAC, including the catalytic triad (S_188_D_342_H_372_) and the oxyanion hole-forming domain (H_110_GGG) (Fig. 7B). The AADAC TMD (M_1_-P_24_) contains three tyrosine residues (Y_16_Y_17_Y_19_) that are highly conserved across species (Fig. 7C). The entry to the catalytic site is protected by an α-helix “lid” (P_29_-L_51_) and the predicted structure showed that the TMD tyrosines Y_16_ and Y_17_ interact with the lid domain (Fig. 7D).

**Fig. 7.**
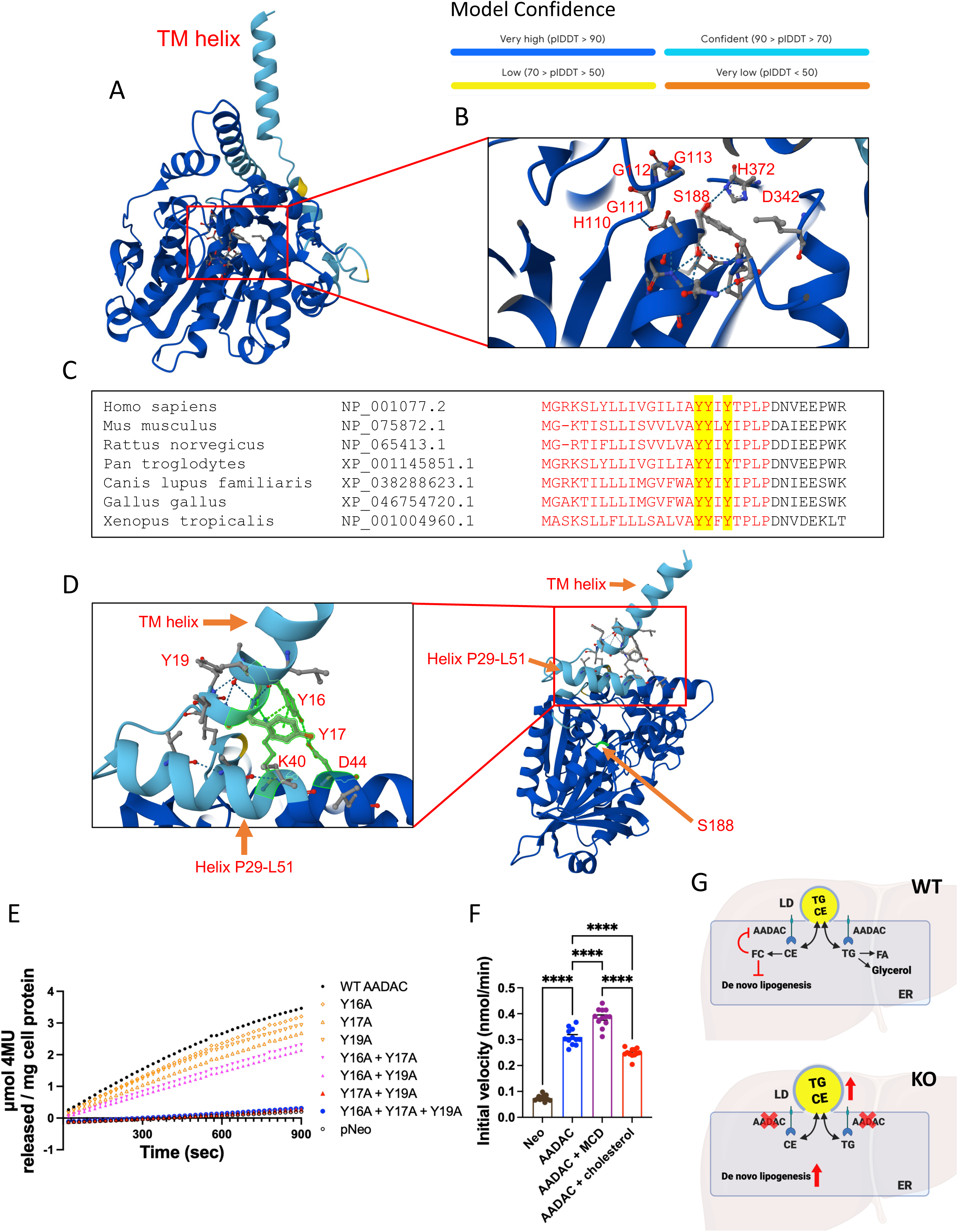
Predicted protein structure of AADAC and regulatory mechanism of AADAC activity. **(A)** Mouse AADAC protein structure predicted by AlphaFold 3. **(B)** Structure of the catalytic site of AADAC. **(C)** Sequence alignment of AADAC transmembrane domain (TMD) across species. TMD regions are shown in red, and the three conserved tyrosine residues are highlighted in yellow. **(D)** Interaction between TM helix and “lid” of AADAC catalytic site. **(E)** Esterase activity of various AADAC mutants transfected in Huh7 cells. **(F)** Esterase activity in McArdle RH-7777 cells stably expressing AADAC after 6-hour treatment with either the cholesterol-sequestering compound methyl-β-cyclodextrin (MCD) or cholesterol loading. Esterase activity is presented as initial velocity from kinetic curves. Data are pooled from three independent experiments and expressed as mean ± SEM. ****P<0.0001. Statistical significance was determined by one-way ANOVA followed by Bonferroni post hoc tests. **(G)** AADAC regulation of hepatic lipid metabolism.

To assess the role of these conserved TMD tyrosines in regulating AADAC activity, we mutated the three tyrosines in various combinations and measured AADAC esterase activity in hepatoma cells. While individual tyrosine-to-alanine substitutions only marginally decreased AADAC activity, mutation of all three tyrosines eliminated the esterase activity. Mutations of Y_16_+Y_17_ or Y_16_+Y_19_ only mildly decreased AADAC activity, but mutation of Y_17_+Y_19_ eliminated AADAC activity to the same degree as the triple mutation (Fig. 7E). Interestingly, immunoblotting showed that mutation of all three tyrosines or Y_17_+Y_19_ decreased AADAC protein abundance, suggesting decreased protein stability (fig. S7).

Tyrosine is an essential residue in conventional cholesterol-binding motifs of membrane proteins and potentially confer affinity for cholesterol (*47*), suggesting a regulatory role of membrane cholesterol binding in AADAC activity. Indeed, depletion of cellular cholesterol by incubation with cholesterol sequestering compound methyl-β-cyclodextrin (MCD) results in enhanced AADAC activity; loading cells with cholesterol results in inhibition of AADAC activity (Fig. 7F).

## Discussion

MASLD, the hepatic consequence of systemic metabolic dysfunction, arises from chronic nutrient excess and disordered lipid synthesis and mobilization in the liver (*31*). However, the pathological progression of MASLD is still not fully understood. Unlike adipocytes, hepatocytes are not specialized for long-term neutral lipid storage. Disruptions in hepatic turnover of preformed lipids contribute directly to MASLD progression but the lipolytic processes responsible for TG and CE hydrolysis in the liver remain incompletely defined (*48*). In this study, we identified AADAC as an important lipase regulating hepatic TG and CE storage. AADAC is predominantly expressed in hepatocytes (*13, 14*). Loss of AADAC from hepatocytes results in decreased neutral lipid turnover and increased TG and CE synthesis culminating in augmented lipid storage in larger size LDs. Importantly, the present study identifies AADAC as a novel hepatic CE hydrolase, as ablation of AADAC leads to CE accumulation in the liver.

Increased lipogenesis is a distinct characteristic of MASLD (*6*). The present study reveals increased *de novo* lipid synthesis regulated by a cascade initiated through the activation of SREBP2 in AADAC deficient livers (Fig. 5F). SREBP2 activity is tightly regulated by changes in ER cholesterol concentration within a narrow range (*5*). As an ER-localized lipase possessing CE hydrolase activity, AADAC may inhibit SREBP2 processing by releasing cholesterol from CE and transiently increasing ER cholesterol concentration. AADAC activity is upregulated by cholesterol depletion and downregulated by cholesterol loading, suggesting a feedback inhibition of its own activity when cholesterol is abundant. Together, these findings suggest a model in which AADAC serves to fine-tune SREBP2 activity and SREBP-dependent *de novo* lipogenesis, thereby restraining excessive lipid synthesis in the postprandial state.

To support enhanced metabolic processes toward lipid synthesis, remodeled TCA cycle and related metabolic pathways, potentially driven by increased SREBP activity, were observed in the hepatic transcriptomic profile of AADAC KO mice. The citrate-malate shuttle transfers acetyl-CoA equivalents from mitochondria to the cytosol, where they serve as building blocks for lipid biosynthesis. Genes encoding key components of the citrate-malate shuttle, *Acly* and *Mdh1*, are established SREBP targets (*40, 42*) that were upregulated in AADAC deficient livers. Consistent with our findings, increased SREBP activity has been reported to promote the diversion of glucose-derived carbons from mitochondria to the cytosol via the citrate-malate shuttle, rather than through the downstream TCA cycle (*49*). In addition to Acly, the other major enzyme for cytosolic acetyl-CoA production and SREBP target, Acss2, was also transcriptionally upregulated in AADAC deficient livers. Elevated ACLY and ACSS2 expression is similarly observed in human MASH (*50*). Dual inhibition of these enzymes reduced MASLD/MASH in various preclinical models (*50, 51*). SREBP also activates *Me1* and the pentose phosphate pathway to support NADPH production required for lipid biosynthesis (*39*). These upregulated pathways, together with enhanced TCA cycle enzyme expression that may lead to increased basal ATP production, could supply both substrates and energy required for elevated *de novo* lipid synthesis in AADAC deficient livers.

AADAC TMD contains a structural triad found in the inverted cholesterol recognition/interaction amino acid consensus (CARC) motif, consisting of a basic lysine residue, central aromatic tyrosine residues, and aliphatic leucine/valine residues (*47*). Protein structure prediction suggests that ER cholesterol may modulate AADAC activity through its interaction with the TMD which is functionally tethered to the ’lid’ helix controlling access to the active site, thereby reducing AADAC activity. The precise structure of cholesterol-AADAC complexes and the conformational changes induced by cholesterol binding or depletion remain to be investigated by structural studies.

A sex-specific pattern in liver lipid profiles between fasting and feeding states after one week of WTD feeding was observed. Fasting increases hepatic neutral lipids in male WT mice but decreases them in female WT mice, which is consistent with previous observations in a human study, as more circulating FAs were directed to peripheral tissues rather than livers in females during fasting (*29*). The increased hepatic neutral lipid levels following refeeding in female WT mice may reflect elevated chylomicron remnant delivery and lipid synthesis in a fed state. AADAC ablation impairs neutral lipid turnover between fasting and refeeding in male mice, resulting in a diminished lipid rebound upon refeeding, while increasing fasting hepatic CE level in female mice in one week of WTD feeding. In the long-term WTD feeding, female AADAC KO mice also showed augmented neutral lipid accumulation in the refed state.

Transcriptome analysis revealed that after one week of WTD, female mouse livers exhibited significantly fewer DEGs between genotypes compared to male mice. Lipid synthesis and metabolism genes that were upregulated in male KO livers showed significantly higher baseline expression in female WT mice compared to male WT mice. This sex-specific difference in lipogenesis and lipogenic gene expression (higher in female than male) has been previously reported in humans and rats following high-fructose diet feeding (*52, 53*). Consistent with this pattern, we observed upregulation of *Mlxipl* (encoding ChREBP) in female WT livers compared to males (annotated to the GO term “fatty acid biosynthetic process” in this comparison). Despite these sex-specific distinctions during short-term feeding, both male and female AADAC KO mice exhibited accelerated development of MASLD features with long-term WTD feeding, suggesting that prolonged nutrient excess ultimately overwhelms these metabolic differences between sexes and reveals the full impact of AADAC deficiency.

Continuous food availability in contemporary lifestyles keeps individuals in a sustained postprandial metabolic state. The feeding state represents a critical window in which efficient hepatic lipid turnover is required to control intrahepatic lipid stores, as adipose tissue lipolysis is suppressed and exogenous fatty acid influx is minimal. In the current study, we demonstrated that AADAC is a lipase crucial for the turnover of preformed neutral lipid in the liver. Additionally, AADAC may restrain *de novo* lipid synthesis by modulating ER cholesterol release from CE through a feedback-regulated mechanism (Fig. 7G), thereby limiting excessive lipid accumulation in postprandial state. Consequently, inactivation of AADAC leads to exacerbated MASLD features in the conditions of chronic overnutrition.

### Methods Mouse Studies

AADAC KO mice (Taconic Biosciences, Model #TF1722) were generated based on a homologous recombination gene-targeting strategy that replaced 4.4 kb, including exons 1 (with the initiation codon) and 2 with a βgeo/puromycin resistance cassette as seen in fig. S1A. Targeting vectors were electroporated into 129/SvEv mouse ES cells and following selection were transferred into C57BL/6 blastocysts. The generated chimeric mice that showed germline transmission were crossed twice with C57BL/6NTac mice to create mixed background lines and then backcrossed a further 8 generations with C57BL/6J mice. Genetic analysis by the Jackson Laboratory confirmed the production of the congenic C57BL/6J background of AADAC KO mice.

All animal procedures were conducted under Animal Use Protocol 402 and in accordance with the Canadian Council on Animal Care guidelines and policies, with approval from the University of Alberta’s Animal Care and Use Committee. Animals were maintained on a 12-hour light/dark cycle (7am-7pm) at 22 ± 2°C and 40-70% humidity, and fed standard chow diet (PICO laboratory Rodent Diet; 5% fat, 0.04% cholesterol). Mice were housed in greenline cages (Tecniplast-GM500) with aspen chip bedding, and the enrichment system in each cage includes a mouse enrichment device (PVC tube), Krinkle paper substrate (8oz standard), and cotton fiber substrate.

For the short-term diet study, 2 - 5 months old male and female wild type (WT) and AADAC KO mice were fed a high-fat, high-cholesterol, high-sucrose western-type diet (WTD, 42% kcal from fat, 0.2% cholesterol, sucrose 34% by weight, Inotiv TD 88137) for one week. Livers and blood were collected from mice at the end of the feeding cycle (7 am, fed), after 12 hours fasting (9pm - 9am, fasting), or after 2 or 6 hours refeeding with WTD following 12 hours fasting. After 2 hours refeeding, intestinal mucosa was collected as previously described (*54*). In brief, small intestines were excised, flushed with PBS containing protease inhibitor cocktail (Sigma), and kept on ice. The small intestine was divided into three portions of length ratio 1:3:2 (corresponding to duodenum:jejunum:ileum). Jejunum sections were fixed in 10% neutral-buffered formalin for histology, or jejunal mucosa was collected after intestinal segments were opened longitudinally and was snap-frozen in liquid nitrogen.

For the long-term diet study, 10 weeks old male and female WT and KO mice were fed WTD for 10 weeks. Tissues were collected after 2 hours refeeding following 12 hours fasting.

For PLIN5 antisense oligonucleotide (ASO) treatment, 12 weeks old WT and AADAC KO male mice were administrated with Plin5 (THA–GalNAc3–conjugated) or control ASO provided by Ionis Pharmaceuticals (Carlsbad, CA) at 40 mg/kg, prepared in sterile saline, and delivered via intraperitoneal injection twice a week for 3 weeks. Mice were then switched from chow diet to WTD for one week while continuing ASO administration using the same procedure. Tissues were harvested in the fed state the next morning after the final ASO dose.

Male *ob/ob* mice homozygous for a spontaneous mutation in the leptin gene (B6.Cg-*Lep^ob^*/J) and the control lean C57BL6/J mice were obtained from the Jackson Laboratory. Liver tissues were collected from 16 weeks old mice fed ad libitum with standard chow diet.

### Histopathological analysis of MASLD

A portion of liver was fixed in 10% neutral buffered formalin, and 5 µm paraffin sections were collected. For MASLD scoring, slides were stained with hematoxylin and eosin, then steatosis was evaluated. The modified NAFLD activity score, used to assess the MASLD progression, was calculated as the sum of steatosis, ballooning, portal inflammation, and lobular inflammation scores obtained from the stained sections (*55, 56*). **Measurement of AADAC esterase activity**

Mouse *Aadac* cDNA, codon optimized for human cells, was synthesized (GenScript Biotech). The cDNA was cloned into the Mlu I and Not I sites of pCI Neo. Site-directed mutagenesis was performed with the QuikChange II site-directed mutagenesis kit (Agilent Technologies) to introduce tyrosine-to-alanine mutations within the transmembrane domain of AADAC. Huh7 cells were transfected with the empty vector pCI-Neo or with the constructs expressing WT or mutated *Aadac* using Lipofectamine 2000 (Thermo) and harvested 24 hours post-transfection.

AADAC stable expressing McA-RH7777 cells were incubated with either 2.5 mM Methyl-β-cyclodextrin (MCD, Sigma C4555) or 100 μM MCD-cholesterol complex (prepared as previously described (*57*)) in DMEM medium. Cells were harvested after 6-hour incubation.

Livers were collected from chow fed WT and AADAC KO mice under fed condition and 20% (w/v) liver homogenate was prepared in sucrose buffer (250 mM sucrose, 50 mM Tris, pH 7.4, 1 mM EDTA). Liver homogenates were spun at 600g at 4°C for 5 minutes to remove unbroken cells, nuclei and debris, the supernatant was centrifugated at 25,000g at 4°C for 10 minutes to pellet mitochondria, peroxisomes and lysosomes, then the 25,000g supernatant was centrifugated at 106,000g at 4°C for 1 hour to pellet microsomes.

Esterase activity was measured utilizing 4-methylumbelliferyl (4-MU) acetate as described previously (*58*). The enzymatic reaction was initiated by the addition of 20 μL of 1mM 4-MU acetate in 20 mM Tris/HCl (pH 8.0), 1 mM EDTA and 300 μM taurodeoxycholate to cell lysates containing 5 μg of protein, or mouse liver homogenates and microsomal fractions containing 3μg of protein in a 96-well plate in a final volume of 200µL. The plate was incubated at 37°C and the release of fluorescent 4-MU was detected with a Fluoroskan Ascent FL Type 374 (Thermo Labsystems) with excitation/emission wavelengths of 355/460nm. Fluorescence values generated with a standard solution of 4-methylumbelliferone (sodium salt) were used to quantify 4-MU release.

### Plasma and tissue chemistries

Enzymatic assays kits were used to measure plasma levels of ketone bodies (WAKO), total cholesterol (WAKO), and triacylglycerol (TG, Roche) according to the manufacturer’s instructions.

The lipid profile was determined in liver homogenates (1 mg of protein) by gas chromatography using tridecanoin as the internal standard as previously described (*59*). In the long-term WTD feeding study, liver lipids were extracted from liver homogenates by the modified Folch method (*60*) and lipid profiles were determined by high-performance liquid chromatography (HPLC) using phosphatidyldimethylethanolamine (PDME) as an internal standard. Liver retinol and total retinyl esters were measured by HPLC following the published method (*61*).

A 10% (w/v) intestinal mucosa homogenate was prepared in sucrose buffer (250 mM sucrose, 50 mM Tris, pH 7.4, 1 mM EDTA) and lipids were extracted by the modified Folch method. TG and TC were determined using Roche and WAKO kits, respectively.

### In vivo VLDL-TG secretion assay

Mice were fasted overnight and then injected intraperitoneally with Poloxamer 407 (1 g/kg body weight). Blood was collected before (“time 0”) and 1, 2, 3, and 4 hours after injection. TG (Roche) and total cholesterol (WAKO) concentrations were determined by kit assays.

For the “refed” VLDL secretion assay, mice were fasted overnight and then injected intraperitoneally with Poloxamer 407 (1 g/kg body weight), followed by oral administration of glucose (2 g/kg body weight). Blood was collected before (“time 0”) and 1, 2, 3, 4 and 5 hours after injection. TG (Roche), ketone bodies (WAKO) and total cholesterol (WAKO) concentrations were determined by kit assays.

### In vivo chylomicron-TG secretion assay

Blood was collected before the experiment as the “time 0”. Fasted mice received an intraperitoneal injection of poloxamer-407 (1 g/kg body weight) and after 30 minutes, mice were gavaged with 150 µl of olive oil containing 10 µCi [^3^H]triolein (Perkin Elmer). Blood was collected at 1, 2, and 4 hours after gavage. Lipids were extracted by modified Folch method (*60*) from plasma and separated by TLC plates with the solvent system heptane:isopropyl ether:acetic acid (15:10:1 by vol). Lipids were visualized by exposure to iodine, and radioactivity in each class of lipid was determined by scintillation counting. **Body composition measurements**

Whole-body lean and fat masses were determined in mice by an EchoMRI™ system after 10 weeks of WTD feeding.

### Glucose tolerance test

Mice were fasted for 16 hours prior to oral gavage of glucose (2 g/kg body weight). Blood glucose concentration was monitored at the indicated time points using glucose strips (Accu-Check glucometer; Roche Diagnostics).

### Insulin tolerance test

Mice were fasted for 6 hours and then injected intraperitoneally with insulin (1 U/kg body weight). Blood glucose concentration was monitored at the indicated time points using glucose strips (Accu-Check glucometer; Roche Diagnostics).

### Blood collection from human subjects

Blood was collected from healthy adult males after overnight fast (fasted) and 4 h after consumption of 235 ml of Ensure Calorie Plus Meal Replacement Drink (Abbott Laboratories) containing 355 calories (28% from fat, 57% from carbohydrate and 15% from protein) following an overnight fast (refed). Blood collection and refeeding protocols were approved by the University of Alberta Human Ethics Board (Pro00059201). All participants provided written informed consent. Sera prepared from the collected blood were pooled and used in experiments as described.

### Detection of lipid droplets (LDs) in primary hepatocytes

Mouse primary hepatocytes were freshly isolated from chow fed 4 months old male AADAC WT and KO mice after 12 hours fasting (9 pm-9 am), and from 2 hours and 6 hours refeeding following 12 hours fasting. Cells were plated on collagen-coated coverslips and incubated with DMEM+15% FBS for 3 hours before proceeding to LD staining with BODIPY 493/503 (Invitrogen) and nuclei staining by DAPI. Coverslips were examined by an Olympus IX-81 CSU10 spinning disk confocal microscope. Images were analyzed by Fiji and at least 30 cells were analyzed for LD morphology.

For LD and mitochondria proximity analysis, primary hepatocytes isolated from WT and KO mice were plated on collagen-coated coverslips and incubated with 200nM MitoTracker® Deep Red FM (Life technologies) for 20 minutes, subsequently fixed by 4% paraformaldehyde, stained with BODIPY 493/503 for LDs and DAPI for nuclei. Images were acquired by an Olympus IX-81 CSU10 spinning disk confocal microscope and analyzed with Fiji. Data are shown as the percentage of LD perimeter in proximity to mitochondria relative to total LD perimeter.

### Hepatocyte double-label pulse-chase experiments to assess cellular lipolysis

Mouse primary hepatocytes were freshly isolated from chow fed 4 months old male AADAC WT and KO mice after 12 hours fasting (9 pm-9 am). 1X10^6^ cells were plated in each 60mm collagen-coated cell culture dish with DMEM medium+10% human fasted serum. To stimulate neutral lipid synthesis, cells were pulse labeled for 4 hours with 2 ml of DMEM containing 10% human fasted serum, 5 µCi [^3^H]oleic acid (OA) (Perkin Elmer)

+ 0.4 mM non-labeled OA bound to 0.5% BSA, 1 µCi [^14^C]glycerol (Perkin Elmer) + 50 µM non-labeled glycerol. Some dishes after the pulse period were harvested to measure incorporation of the labeled precursors into lipids. Other dishes were changed to a chase medium containing low glucose (1 g/L) DMEM, 0.5% BSA, 5 µM DGAT1 and DGAT2 inhibitors (PF-04620110 and PF-06424439, respectively) synthesized by Pfizer Inc. (*62, 63*) and purchased from Sigma-Aldrich, 50 nM insulin, with or without 200 µM diethyl-p-nitrophenyl phosphate (E600, pan-lipase inhibitor), and incubated for 12 hours. Medium was collected, lipids were extracted with the modified Folch method (*60*) after acidification allowing partitioning of unesterified FA into the organic (lipid) phase and [^3^H] radioactivity in the aqueous phase was counted using a liquid scintillation counter to determine the release of acid-soluble metabolites from cells. Cells were harvested in PBS, and lipids extracted from cell lysates were separated by thin layer chromatography (TLC) with the solvent system heptane:isopropyl ether:acetic acid (15:10:1 by vol). Lipids were visualized by exposure to iodine, and radioactivity in TG and CE was determined by liquid scintillation counting. The percentage of lipid turnover in the chase phase was expressed relative to its level in pulse cells.

### Hepatocyte metabolic labeling study to assess *de novo* lipogenesis

Mouse primary hepatocytes were freshly isolated from 4 months old male AADAC WT and KO mice fed with chow diet after 5-6 hours refeeding preceded by 12 hours fasting. 1X10^6^ cells were plated in each 60mm collagen-coated cell culture dish with DMEM medium + 10% serum prepared from refed human individuals. To assess hepatic *de novo* lipogenesis, hepatocytes were incubated for 4 hours in 2 ml DMEM containing 10% human refed serum, 10 μCi [^3^H]acetic acid (Perkin Elmer) and 50 μM non-radiolabeled acetic acid. Cells were harvested, lipids were extracted, resolved by TLC, and radioactivity in various lipid classes was measured.

### Electron Microscopy

Mouse livers were harvested after 1 week of WTD feeding. Tissues were immersed into conventional TEM fixative (3% glutaraldehyde and 3% paraformaldehyde in 0.1M sodium cacodylate buffer) (w/v =1/5) for an hour for the initial fixation. Samples were then dissected into 1-2 mm^2^ pieces and transferred to glass vials filled with ice cold conventional TEM fixative (w/v=1/10). Samples were post-fixed in 1% potassium ferrocyanide and 1% osmium tetroxide in 0.1M sodium cacodylate buffer, dehydrated in ethanol, and embedded in Spurr’s resin. Sections were obtained by Leica EM UC7 Ultramicrotome and imaged with Hitachi H-7650 Transmission Electron Microscope.

### Microarray

Intestinal mucosa was collected from male WT and AADAC KO mice in the 12-hour fasted or 2-hour refed states after one week of WTD feeding (n=5-6 per group, 23 samples in total). Total RNA was extracted using Trizol (Invitrogen) and RNA integrity was confirmed using Agilent 2100 Bioanalyzer RNA Nanochips (Agilent Technologies). Microarray analysis was performed at The Centre for Applied Genomics (Toronto, ON, Canada) using the Affymetrix Mouse Gene 2.0 ST array. Raw intensity values were pre-processed using the rma() function from the oligo package, which implements Robust Multi-array Average (RMA) background correction, quantile normalization, and probe summarization into transcripts via median polish. Control probes were removed before downstream analyses. Probe annotation was based on the mouse reference genome mm10. Differential expression (DE) analysis was performed using limma, and statistical significance was determined using a Benjamini-Hochberg adjusted p-value (FDR) threshold of < 0.05. Gene Ontology (GO) enrichment and KEGG pathway analyses were performed using clusterProfiler (version 4.14.4) in R with an FDR threshold of < 0.1.

### RNA sequencing (RNA-seq)

Liver samples were collected from 10 weeks old male and female WT and AADAC KO mice (n=5-6 per group, 23 samples in total) in the fed state after one week of WTD feeding. Total RNA was extracted using Trizol (Invitrogen) and cleaned up with a RNeasy Mini Kit (Qiagen). RNA integrity was confirmed using Agilent 2100 Bioanalyzer RNA Nanochips (Agilent Technologies). Library preparation and sequencing were performed at Canada’s Michael Smith Genome Sciences Centre (Vancouver, BC, Canada). In brief, First-strand cDNA was synthesized from heat-denatured purified mRNA using a Maxima H Minus First Strand cDNA Synthesis kit (Thermo) with random hexamer primers, followed by bead purification using PCR Clean DX (Aline Biosciences) on a Microlab NIMBUS robot (Hamilton Robotics). Second-strand cDNA was synthesized following the NEBNext Ultra Directional Second Strand cDNA Synthesis protocol (NEB) with dUTP incorporation. cDNA was fragmented to 250-300 bp using a Covaris LE220 instrument, followed by end-repair, A-tailing, and Illumina adapter ligation. Products were digested with USER enzyme and amplified with 10 cycles of indexed PCR using NEBNext Ultra II Q5 polymerase (NEB) and Illumina’s dual index primer set. Final libraries were purified and size-selected with PCR Clean DX beads, with quality control performed using Agilent DNA 1000 assay and Qubit quantification before pooling for paired-end 150 bp sequencing on the Illumina NovaSeq X system. Raw sequencing reads were aligned to the mouse reference genome mm10 using STAR (version 2.5.2b), gene and isoform expression were quantified using RSEM (version 1.3.0) with Ensembl 100 gene annotation.

DE analysis was performed using R (version 4.4.1) and DESeq2 package (version 1.46.0) (*64*) with expected read counts (gene-level abundances) as input data. Statistical significance was determined using a Benjamini-Hochberg adjusted p-value (FDR) threshold of < 0.05. GO enrichment and KEGG pathway analyses were performed using clusterProfiler (version 4.14.4) in R with an FDR threshold of < 0.05. RNA-seq analysis results were visualized using ggplot2 (version 3.5.1), pheatmap (version 1.0.12), and GOplot (version 1.0.2) in R. STRING network analysis was performed by Cytoscape (Version: 3.10.3) with the confidence cutoff 4.0, followed by Markov Cluster Algorithm (MCL) clustering (Granularity parameter 4) using STRING confidence scores as edge weights.

### Seahorse XF Mitochondrial Stress Test

Mice were fed either chow diet or WTD for one week. Primary hepatocytes were isolated from fed state mice and seeded at a density of 4,500 cells per well in XF96 cell culture microplates (Agilent Technologies) coated with 50 μg/mL collagen I solution (Enzo). Cells were cultured overnight in DMEM containing 15% FBS at 37°C with 5% CO₂. Prior to analysis, culture medium was replaced with unbuffered XF base medium (Agilent Technologies) supplemented with 10 mM glucose, 1 mM pyruvate, and 2 mM glutamine, and cells were equilibrated for 1 hour at 37°C in a non-CO₂ incubator. Mitochondrial bioenergetics were assessed using the Seahorse XF96 Analyzer (Agilent Technologies) with sequential injection of oligomycin (2 μM), carbonyl cyanide 4-(trifluoromethoxy)phenylhydrazone (FCCP, 1.5 μM), and rotenone plus antimycin A (1.5 μM each). Oxygen consumption rate (OCR) was measured for each condition with 3-minute mixing and 3-minute measurement periods. Then ATP-linked respiration was calculated as the difference between basal OCR and oligomycin-inhibited OCR. Maximal respiration was determined as the peak OCR following FCCP injection. Spare respiratory capacity was calculated as the difference between maximal and basal respiration. Following Seahorse analysis, cells were stained with Hoechst 33342 (20 μM) for nuclei visualization and cell counting. All OCR values were normalized to the number of nuclei per well.

### Immunoblot analyses

Livers were homogenized in sucrose buffer (250 mM sucrose, 50 mM Tris, pH 7.4, 1 mM EDTA), homogenate proteins were resolved by SDS-polyacrylamide gels and transferred to PVDF membranes (catalog #IPVH00010; Millipore). Antibodies used in this study include anti-AADAC (1:5,000 dilution, rabbit polyclonal antibody generated in our laboratory), anti-adipose triglyceride lipase (ATGL) (1:1,000 dilution, catalog #2138; Cell Signaling), anti-calnexin (1:5,000 dilution, catalog # SPA-865; Stressgen), anti-GAPDH (1:5,000 dilution, catalog #ab8245; Abcam), anti-perilipin 2 (PLIN2) (1:1,000 dilution, catalog # ab108323; Abcam), anti-perilipin 5 (PLIN5) (1:2,000 dilution, catalog #GP31; PROGEN), anti-VDAC1 (1:1,000 dilution, catalog # ab14734, Abcam), anti-OXPHOS cocktail (1:1,000 dilution, catalog #ms604, MitoScience). Immunoreactivity was detected by enhanced chemiluminescence and visualized by G:BOX system (SynGene). The relative intensities of the resulting bands on the blots were analyzed by densitometry using the GeneTools program (SynGene).

### RNA isolation and real-time qPCR analysis

Total liver RNA was isolated using Trizol reagent (Invitrogen). First-strand cDNA was synthesized from 2 μg total RNA using Superscript ΙΙΙ reverse transcriptase (Invitrogen) primed by oligo (dt)_12-18_ (Invitrogen) and random primers (Invitrogen). Real-time qPCR was performed with Power SYBR^®^ Green PCR Master Mix (Life Technologies) using the StepOnePlus-Real time PCR System (Applied Biosystems). Data were analyzed with the StepOne software (Applied Biosystems). Standard curves were used to calculate mRNA abundance relative to that of a control gene encoding cyclophilin (*Ppia*). Real-time qPCR primers are summarized in the table S3. All primers were synthesized by Integrated DNA Technologies.

### Statistical analysis

Data are presented as mean ± SEM. Differences among group means were assessed by two-way ANOVA followed by Bonferroni post hoc tests, and the unpaired *t-* test was used for two-group comparisons (GraphPad PRISM 10 software). Statistical significance was defined as *P < 0.05, **P < 0.01, ***P < 0.001, ****P < 0.0001.

### Data availability

Microarray data have been deposited in the Gene Expression Omnibus (GEO) database under accession number GSE318918. RNA-seq data have been deposited in the GEO database under accession number GSE312618.

## Acknowledgements

Lipid analysis was performed at the University of Alberta Faculty of Medicine & Dentistry Lipidomics Core, and imaging was performed at the University of Alberta Faculty of Medicine & Dentistry Cell Imaging Centre. Both facilities receive financial support from the Faculty of Medicine & Dentistry and Canada Foundation for Innovation (CFI) awards to contributing investigators. We thank Dr. Chinmayee Das for assistance with confocal image acquisition and quantification for the peridroplet mitochondria analysis.

