## Supplemental Figures for "Endoplasmic Reticulum Associated Lipolysis Regulates Hepatic Fat Synthesis and Turnover"

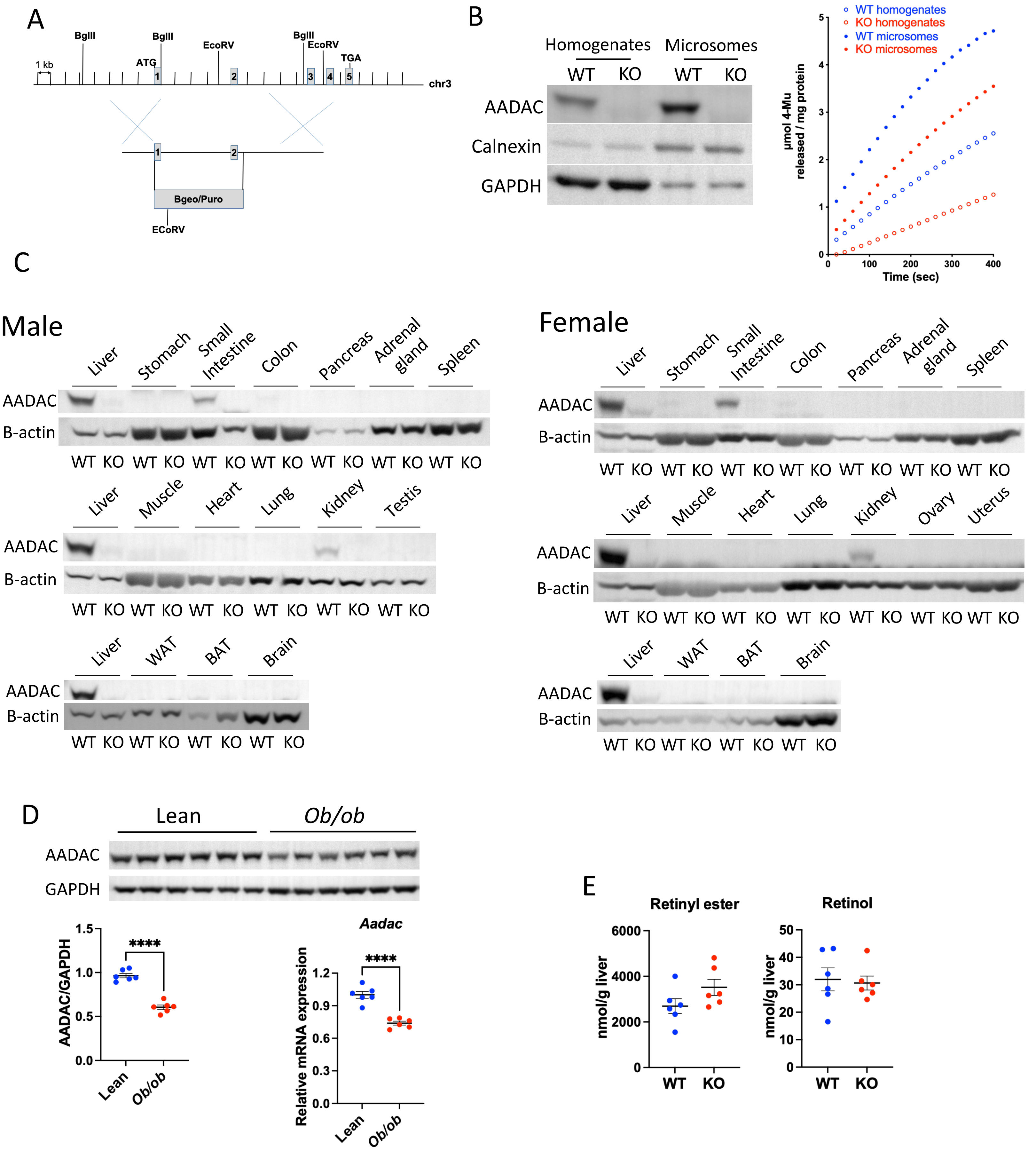


**Fig. S1. Generation of AADAC KO mice**. **(A)** Strategy of AADAC KO mice generation. **(B)** Immunoblotting and activity of AADAC in liver homogenate and microsomal fractions prepared from WT and AADAC KO mice. ER marker calnexin, and cytosolic marker GAPDH were used as loading controls. **(C)** Tissue distribution of AADAC in mice analyzed by immunoblotting. Equal amount of tissue homogenate protein was analyzed. Liver samples were included in every blot for comparison. **(D)** Decreased AADAC protein and *Aadac* mRNA abundance in the livers of *ob/ob* mice. **(E)** Total liver retinyl ester and retinol concentrations in male WT and AADAC KO mice fed with chow diet after 5-7 hours refeeding. Data are expressed as mean ± SEM. ****P<0.0001. Statistical significance was determined by unpaired two-tailed *t*-tests.

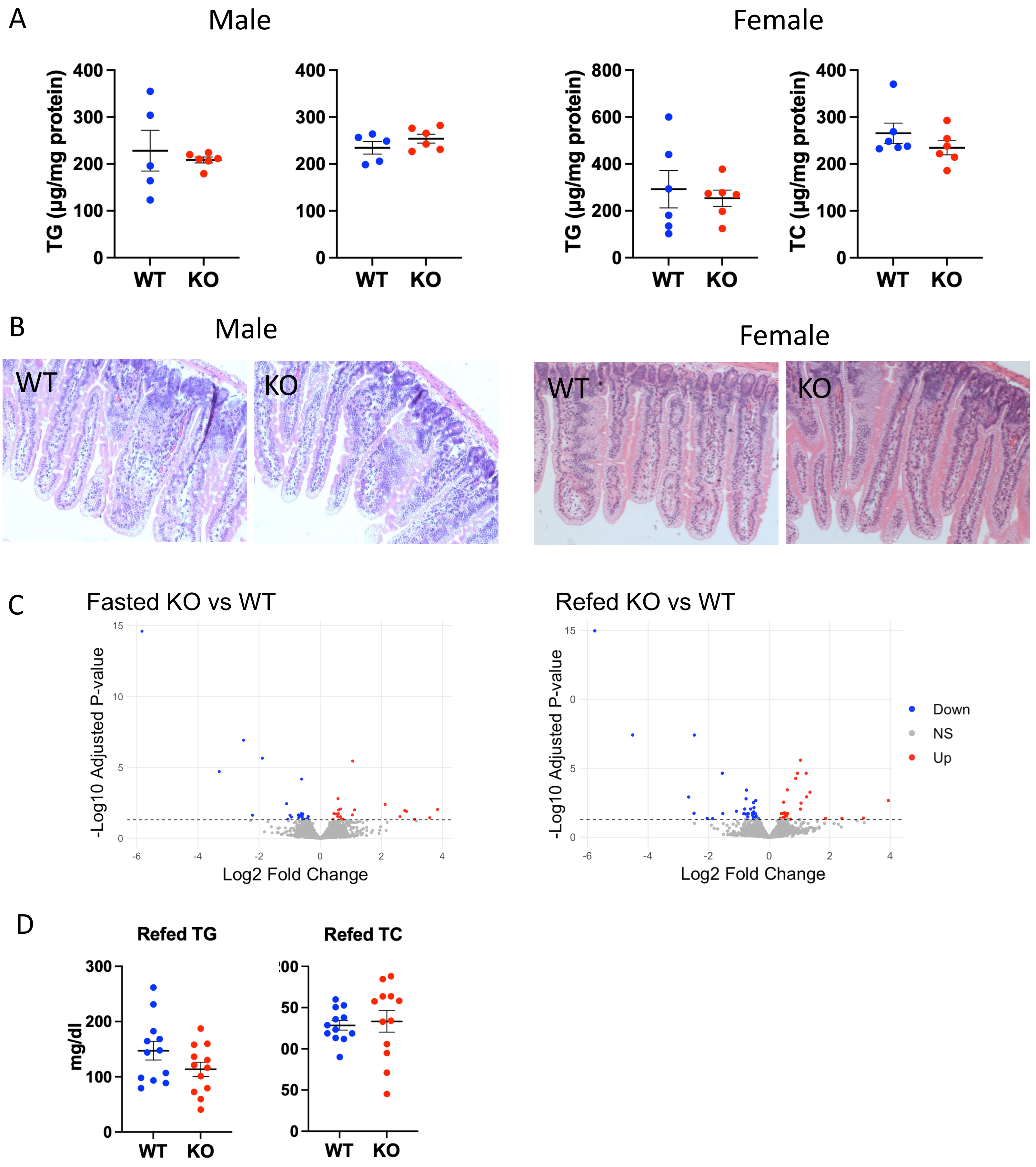


**Fig. S2. AADAC deficiency does not alter intestinal lipid metabolism. (A)** Intestinal mucosal lipid levels and **(B)** histology in the jejunum of WT and AADAC KO mice after 2-hour refeeding following one week of WTD feeding. **(C)** Volcano plot of gene expression in intestinal mucosa of WT and AADAC KO mice assessed by microarray after fasting or 2-hour refeeding following one week of WTD feeding. **(D)** Plasma TG and TC concentrations after 2-hour refeeding in male WT and AADAC KO mice fed with WTD for one week. Data are expressed as mean ± SEM. Statistical significance was determined by unpaired two-tailed *t*-tests.

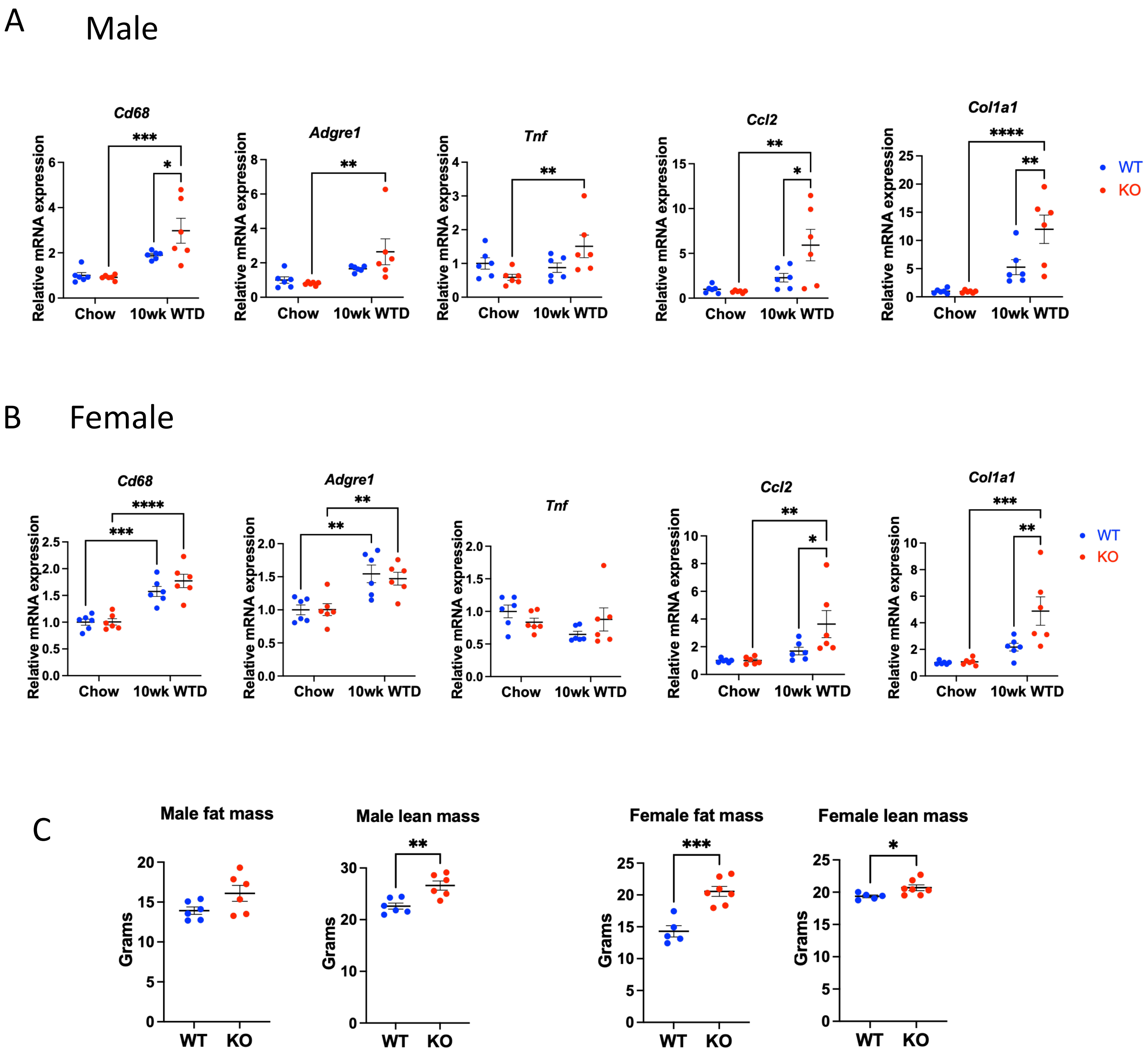


**Fig. S3. AADAC deficiency exacerbates WTD-induced MASLD development and affects whole body metabolism.** QPCR analysis of hepatic expression of inflammation and fibrosis genes in chow and 10 weeks WTD fed **(A)** male and **(B)** female WT and AADAC KO mice. *Cd68*, encoding cluster of differentiation 68; *Adgre1*, encoding F4/80; *Tnf*, encoding tumor necrosis factor alpha; *Ccl2*, encoding monocyte chemotactic protein-1; *Col1a1*, encoding collagen type I alpha 1. **(C)** Absolute fat and lean mass weight of WT and AADAC KO mice fed WTD for 10 weeks. Data are expressed as mean ± SEM. *P<0.05, **P<0.01, ***P<0.001, ****P<0.0001. Statistical significance was determined by two-way ANOVA followed by Bonferroni post hoc tests, or by unpaired two-tailed *t*-tests for two-group comparisons.

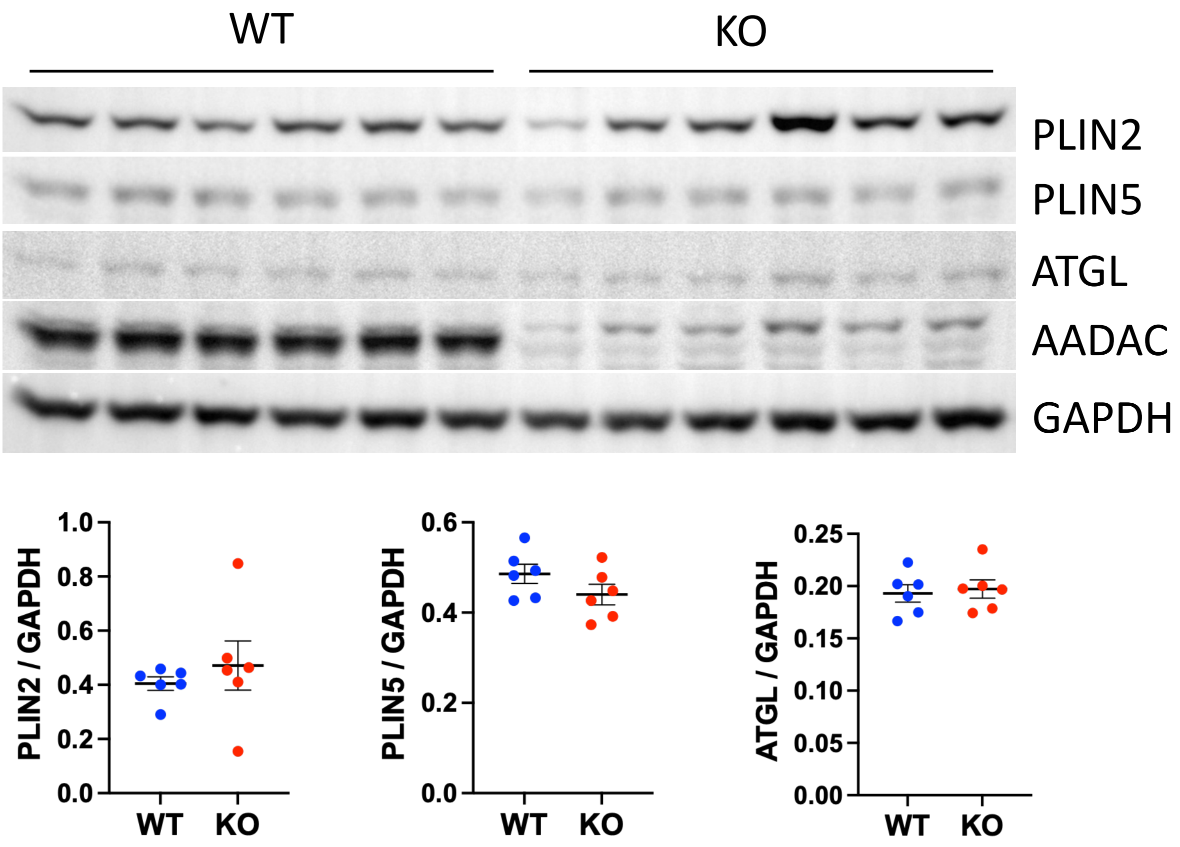


**Fig. S4. Liver PLIN2, PLIN5 and ATGL protein abundance in female mice in the fed state after one week of WTD.** Data are expressed as mean ± SEM. Statistical significance was determined by unpaired two-tailed *t*-tests.

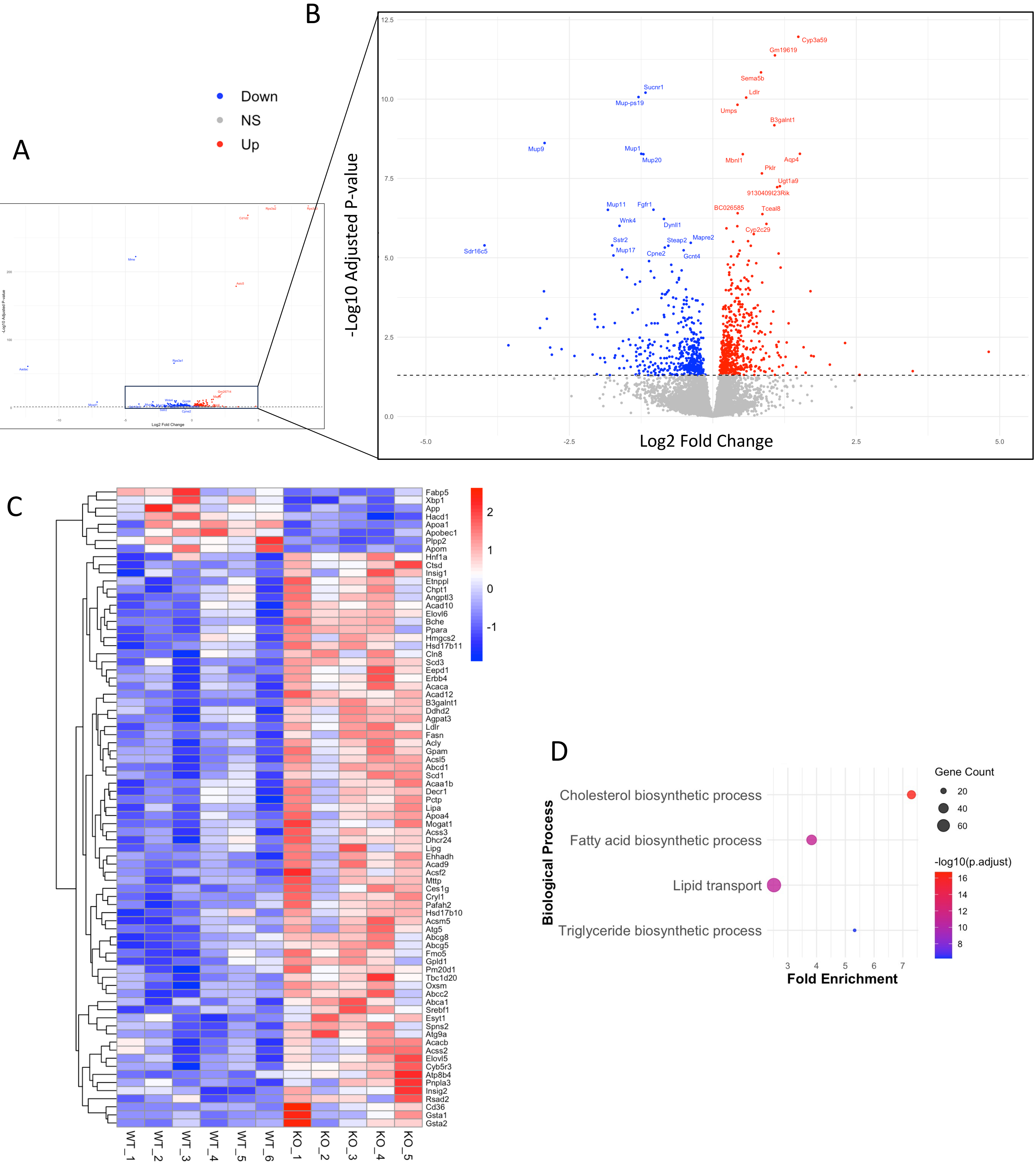


**Fig. S5. RNA sequencing analysis in the livers of AADAC WT and KO mice in the fed state after one week of WTD feeding. (A)** Volcano plot of gene expression changes in the livers of male AADAC KO mice compared to WT mice. Top 20 most upregulated and downregulated genes ranked by adjusted p-value were labelled on the plot. **(B)** A zoomed-in view of the central portion of the volcano plot for better visualization. **(C)** Heatmap of differentially expressed genes (DEGs) related to lipid metabolism in the livers of male AADAC KO mice compared to WT mice. **(D)** Upregulated biological processes in the livers of female WT mice versus male WT mice after one week of WTD feeding.

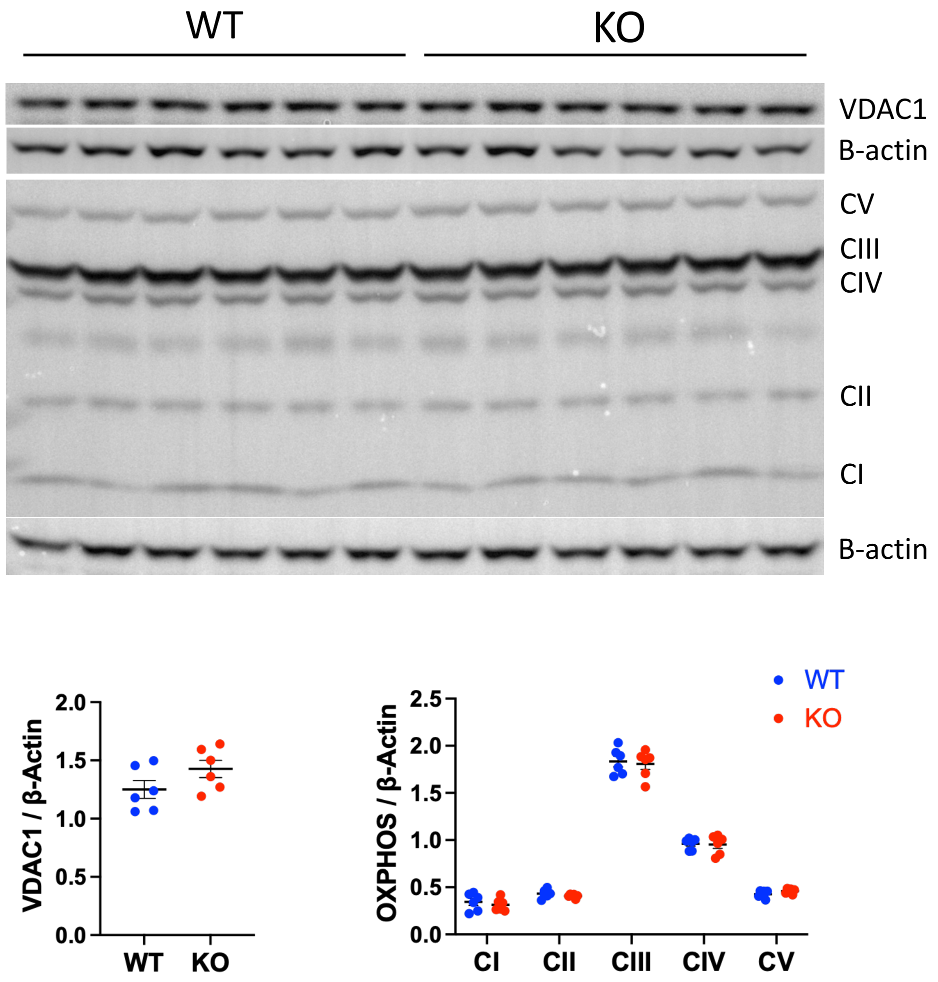


**Fig. S6. Immunoblotting of mitochondrial marker VDAC1 and oxidative phosphorylation (OXPHOS) complexes in the livers of male WT and AADAC KO mice isolated at fed state after one week of WTD.** Data are expressed as mean ± SEM. Statistical significance was determined by unpaired two-tailed *t*-tests.

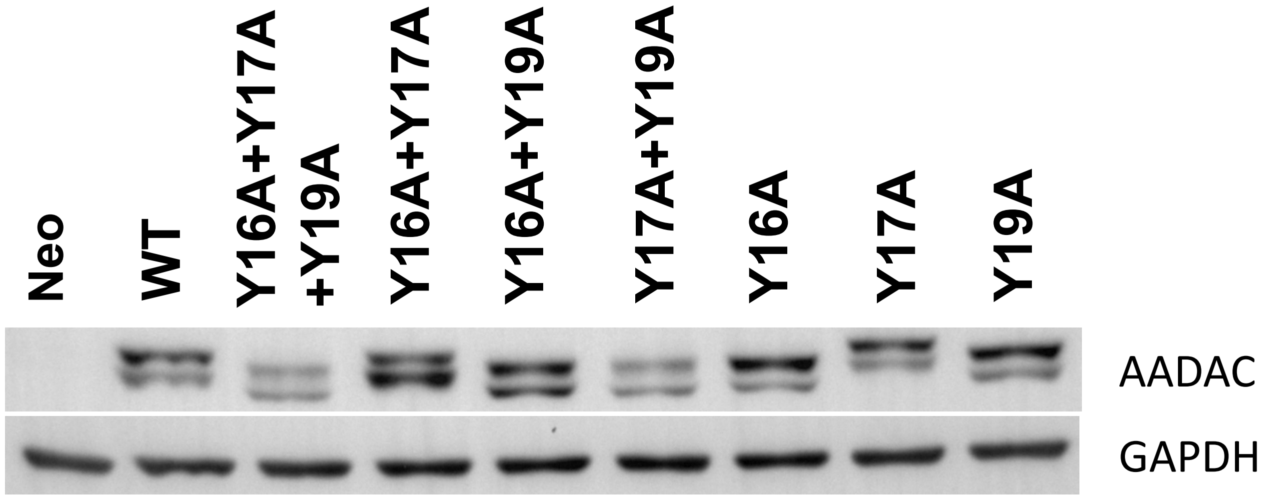


**Fig. S7. AADAC protein abundance of various mutants expressed in the Huh7 cells. GAPDH was used as the loading control.**
