## Supplementary material for "Endoplasmic Reticulum Associated Lipolysis Regulates Hepatic Fat Synthesis and Turnover": Table S3

Table S3. Primers used for quantitative PCR (qPCR) analysis

| **Gene** | **Sequence** |
| --- | --- |
| *Aadac* | F: 5’- ATA TAC CGC TTC CAG ATG CTA TT -3’ |
|  | R: 5’- TAT ATG CGG ACG GGA ACA CT -3’ |
| *Cd68* | F: 5’- GCG GCT CCC TGT GTG TCT GAT -3’ |
|  | R: 5’- GGG CCT GTG GCT GGT CGT AG -3’ |
| *Col1a1* | F: 5’- AGA CAT GTT CAG CTT TGT GGA C -3’ |
|  | R: 5’- GCA GCT GAT TTC AGG GAT G -3’ |
| *Adgre1* | F: 5’- CCC TCG GGC TGT GAG ATT GTG -3’ |
|  | R: 5’- TGG CCA AGG CAA GAC ATA CCA G -3’ |
| *Ccl2* | F: 5’- CAT CCA CGT GTT GGC TCA -3’ |
|  | R: 5’- GAT CAT CTT GCT GGT GAA TGA GT -3’ |
| *Ppia* | F: 5' - TCC AAA GAC AGC AGA AAA CTT TCG - 3' |
|  | R: 5' - TCT TCT TGC TGG TCT TGC CAT TCC - 3' |
| *Tnf* | F: 5’- GTC TAC TGA ACT TCG GGG TGA -3’ |
|  | R: 5’- CAC CAC TTG GTG GTT TGC TAC GAC -3’ |
